# Improving representations of genomic sequence motifs in convolutional networks with exponential activations

**DOI:** 10.1101/2020.06.14.150706

**Authors:** Peter K. Koo, Matt Ploenzke

## Abstract

Deep convolutional neural networks (CNNs) trained on regulatory genomic sequences tend to build representations in a distributed manner, making it a challenge to extract learned features that are biologically meaningful, such as sequence motifs. Here we perform a comprehensive analysis on synthetic sequences to investigate the role that CNN activations have on model interpretability. We show that employing an exponential activation to first layer filters consistently leads to interpretable and robust representations of motifs compared to other commonly used activations. Strikingly, we demonstrate that CNNs with better test performance do not necessarily imply more interpretable representations with attribution methods. We find that CNNs with exponential activations significantly improve the efficacy of recovering biologically meaningful representations with attribution methods. We demonstrate these results generalise to real DNA sequences across several *in vivo* datasets. Together, this work demonstrates how a small modification to existing CNNs, i.e. setting exponential activations in the first layer, can significantly improve the robustness and interpretabilty of learned representations directly in convolutional filters and indirectly with attribution methods.

## Introduction

Convolutional neural networks (CNNs) applied to genomic sequence data have become increasingly popular in recent years, demonstrating state-of-the-art accuracy on a wide variety of regulatory genomic prediction tasks^1–4^, including transcription factor (TF) binding and chromatin accessibility. These successes have been attributed to a CNN’s ability to automatically learn features directly from the training data and make accurate predictions in an end-to-end fashion^5^. However, CNNs, and deep learning models more broadly, have a reputation of being black boxes, with little understanding of their inner workings.

Recent progress to understand model predictions has been driven by attribution methods – such as saliency maps^6^, integrated gradients^7^, DeepLIFT^8^, DeepSHAP^9^, and in genomics, *in silico* mutagenesis^2, 10^ – and other interpretability methods – including Grad-CAM^11^, enhanced integrated gradients^12^, and class optimization of the inputs^6, 13–15^, which has recently been utilized for sequence design^4, 16, 17^, among other existing methods that have not yet been explored thoroughly in genomics^18–20^. Attribution methods are of special interest in genomics because they provide the independent contribution of each input nucleotide toward model predictions – a technique that naturally extends itself to scoring the functional impact of genomic variation, such as single nucleotide polymorphisms. In practice, the attribution “maps” can be challenging to interpret, requiring downstream analysis to obtain more interpretable features, such as sequence motifs, by averaging clusters of attribution scores^21^. In computer vision, it has been demonstrated that there is no guarantee that attribution methods will reveal features that are human interpretable, even if the CNN is capable of a high classification performance^22–24^. Understanding the properties that influence a CNN’s interpretability with attribution methods remains an open problem.

In genomics, an alternative approach to gain insights from a trained CNN is to visualize first layer filters, which requires minimal posthoc analysis to obtain representations of “salient” features, such as sequence motifs. However, it was recently shown that design choices can significantly affect the extent that filters learn motif representations^25, 26^. For instance, pre-convolution weight transformations that model the first layer filters as position weight matrices (PWMs) may be used to learn sequence motifs through the weights^26^. Another CNN design choice employs a large max-pool window size after the first layer, which obfuscates the spatial ordering of partial features, preventing deeper layers from hierarchically assembling them into whole feature representations^25^. Hence, the CNN’s first layer filters must learn whole features, because it only has one opportunity to do so. One drawback to these design principles is that they are limited to shallower networks. Depth of a network significantly increases its expressivity^27^, enabling it to learn a wider repertoire of features. In genomics, deeper networks have found greater success at classification performance^3, 4, 28^. Evidently, there seems to be a trade off between performance and interpretability that goes hand-in-hand with network depth.

One consideration for CNN filter interpretability that has not been comprehensively explored thus far is the activation function. Here we perform systematic experiments on synthetic data that recapitulates a multi-class classification task to explore how first layer activations affect representation learning of sequence motifs. We find the extent that first layer filters learn motif representations is highly dependent on CNN design choice for common activation functions. Strikingly, we find that an exponential activation, which to the authors’ knowledge has never been applied to hidden layers of CNNs, consistently results in robust motif representations irrespective of the network’s depth. We then investigate how CNN design choice influences the efficacy of recovering meaningful representations with attribution methods. We find that CNNs that make more accurate predictions on held-out test sequences do not necessarily recover biologically meaningful representations with attribution methods. One consistent trend that emerges from this study is CNNs that learn robust representations of sequence motifs in first layer filters significantly improve the efficacy of attribution methods. We demonstrate that these results generalize to real DNA sequences across several *in vivo* datasets.

### Exponential activations lead to interpretable motifs

The rectified linear unit (relu) is the most commonly employed CNN activation function in genomics^29^. Alternative activations include sigmoid, tanh, softplus^30^, and the exponential linear unit (elu)^31^ (Table 1). Many common activation functions scale linearly for positive inputs, with differences arising from how they deal with negative inputs (Fig. 1a). Unlike previous activations, we are intrigued by the exponential activation, because it provides a function that is bounded by zero for negative values and diverges quickly to infinity for positive values. Unlike relu or softplus activations, which also bound negative values to zero but scale positive values linearly, the highly divergent exponential function provides a high sensitivity which, in principle, can amplify positive signal while maintaining low background levels. The inputs to the exponential function should be scaled to the sensitive region of the function – optimal scaling varies with the signal and background levels. By setting the activation to be a standard exponential function (Table 1), the network can choose its own threshold by scaling pre-activations with first layer filters. Moreover, the linear behavior of relu and softplus activations can be more permissive in the sense that if background is propagated through the first layer, then deeper layers can still build representations that correct for this noise. On the other hand, for a CNN with exponential activations in the first layer and relu activations in deeper layers, if background noise is propagated through the first layer, then the rest of the network, which is scaled linearly, is ill-equipped to deal with such exponentially amplified noise. In this scenario, we anticipate a failure of training, which would be realized as poor classification accuracy. To be successful, the network must opt for a strategy to suppress background prior to activation and only propagate discriminatory signals, which we anticipate will lead to more interpretable first layer filters. For these reasons, we propose that the exponential activation should only be applied to a single layer of a deep CNN – the layer desired to have interpretable parameters, while employing traditional activations, such as a relu, for the other layers. For genomics, motif representations in first layer filters is highly desirable and hence is the ideal layer for exponential activations. In other applications, determining which layer should employ the exponential activation requires prior knowledge of the relevant scales of the features that are important. To the authors’ knowledge, the exponential activation has not been used as an activation function in hidden layers of CNNs.

**Table 1.**
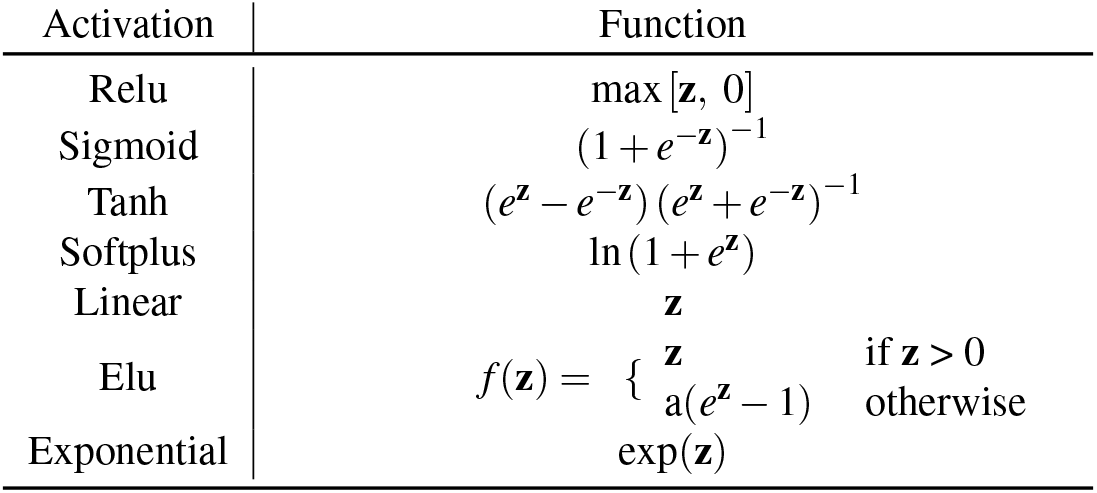
Standard activation functions. This table shows the activation functions for pre-activation values, **z**, in a hidden layer.

| Activation | Function |
| --- | --- |
| Relu | $\max[\mathbf{z}, 0]$ |
| Sigmoid | $(1 + e^{-\mathbf{z}})^{-1}$ |
| Tanh | $(e^{\mathbf{z}} - e^{-\mathbf{z}}) (e^{\mathbf{z}} + e^{-\mathbf{z}})^{-1}$ |
| Softplus | $\ln(1 + e^{\mathbf{z}})$ |
| Linear | $\mathbf{z}$ |
| Elu | $f(\mathbf{z}) = \begin{cases} \mathbf{z} & \text{if } \mathbf{z} > 0 \\ a(e^{\mathbf{z}} - 1) & \text{otherwise} \end{cases}$ |
| Exponential | $\exp(\mathbf{z})$ |

**Figure 1.**
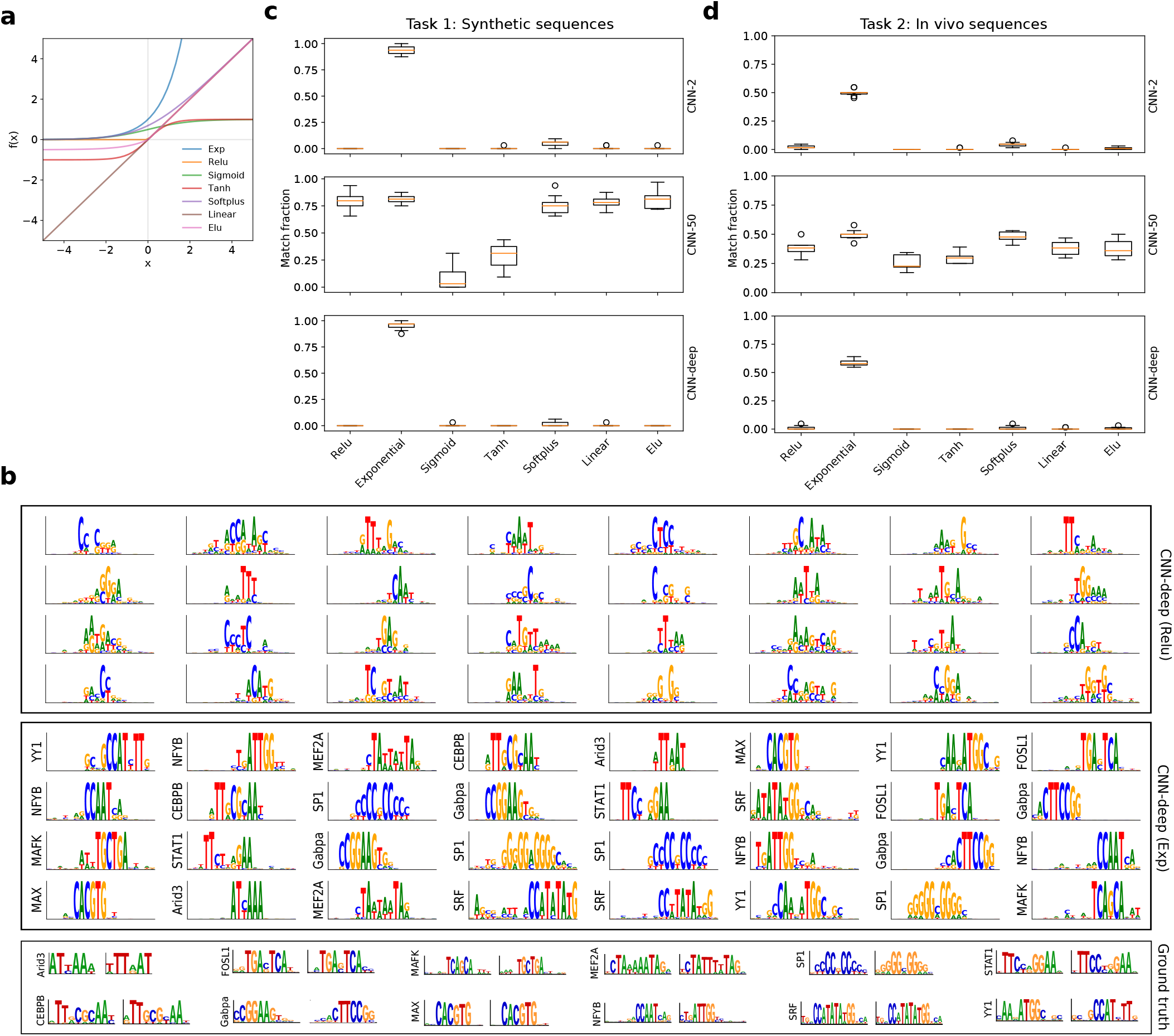
Motif representation performance. (**a**) Plot of various activation functions, including exponential (exp), relu, sigmoid, tanh, softplus, linear, and elu. (**b**) Sequence logos for first convolutional layer filters are shown for CNN-deep with relu activations (top) and exponential activations (middle). The sequence logo of the ground truth motifs and its reverse-complement for each transcription factor is shown at the bottom. The *y*-axis label on select filters represent a statistically significant match to a ground truth motif as determined by Tomtom with an E-value threshold of 0.1. None of the filters from CNN-deep with relu activations yield any hits to ground truth motifs. (**c**) Boxplot of the fraction of filters that match ground truth motifs for CNN-2 (top), CNN-50 (middle), and CNN-deep (bottom) with various first layer activations trained on synthetic sequences of Task 1. (**d**) Boxplot of the fraction of filters that match ground truth motifs for CNN-2 (top), CNN-50 (middle), and CNN-deep (bottom) with various first layer activations trained on real DNA sequences of Task 2. (**c**-**d**) Each boxplot represents the performance across 10 models with different random intializations (box represents first and third quartile and the red line represents the median).

To test the extent that CNN activations influence representation learning, we uniformly trained and tested various CNNs with different first layer activation functions on a multitask classification dataset from Ref.^25^, which we refer to as Task 1. The dataset in Task 1 consists of synthetic DNA sequences embedded with motifs randomly chosen with replacement from a bag of 12 TF motifs from the JASPAR database^32^. Each TF motif represents a unique class. The goal is to determine class membership based on the presence of motifs in each sequence. We explored 3 CNNs, namely CNN-2, CNN-50, and CNN-deep (see Methods for network details and training procedure). Using representation learning design concepts developed for CNNs that employ relu activations^25^, CNN-50 and CNN-2 employ 2 convolutional layers with max-pooling after each convolutional layer, followed by a fully-connected hidden layer. CNN-50 is designed with large max-pooling after the first convolutional layer, which provides an inductive bias to learn “local” representations (*i.e.* whole motifs) in first layer filters, while CNN-2 employs small max-pooling, which allows it to build “distributed” representations in a hierarchical manner, combining partial motifs learned in the first layer into a whole motif representation in deeper layers. CNN-deep consists of 4 convolutional layers with small max-pooling sizes followed by a fully connected layer, which like CNN-2, should build distributed representations of motifs.

The classification performance as measured by the area under the precision-recall curve (AUPR) on a held out test set for Task 1 are more-or-less comparable for each combination of network and activation (Table 2), with CNN-deep exhibiting a slight overall edge. CNN-50 systematically yields a slightly lower performance, with tanh and sigmoid activations yielding the poorest classification accuracy. Tweaking the initialization strategy could presumably improve the classification performance^33^, but was not explored here to maintain a systematic approach. Although the unbounded behavior of the exponential could make the CNN activations diverge, in practice, there were no obvious issues with training (Supplemental Fig. 1), with convergence times that are similar to CNNs with relu activations and stable gradients throughout (Supplemental Fig. 2).

**Table 2.** Filter performance comparison. This table shows the average area under the precision-recall curve (AUPR) across the 12 TF classes, average percent match between the first layer filters and the entire JASPAR vertebrates database (JASPAR), and the average percent match to any ground truth TF motif (Relevant) for different CNNs for Task 1 (Synthetic) and Task 2 (*In vivo*). The errors represent the standard deviation of the mean across 10 independent trials using random intializations.

| MODEL | ACTIVATION | CLASSIFICATION<br>AUPR | SYNTHETIC |  | CLASSIFICATION<br>AUPR | *In vivo* |  |
| --- | --- | --- | --- | --- | --- | --- | --- |
|  |  |  | MOTIF MATCH<br>(JASPAR) | MOTIF MATCH<br>(RELEVANT) |  | MOTIF MATCH<br>(JASPAR) | MOTIF MATCH<br>(RELEVANT) |
| CNN-DEEP | RELU | 0.898±0.002 | 0.266±0.078 | 0.000±0.000 | 0.675±0.003 | 0.259±0.029 | 0.011±0.016 |
|  | EXP | 0.906±0.002 | 0.953±0.038 | 0.953±0.038 | 0.669±0.002 | 0.859±0.038 | 0.584±0.030 |
|  | SIGMOID | 0.892±0.020 | 0.106±0.054 | 0.003±0.009 | 0.667±0.003 | 0.078±0.020 | 0.000±0.000 |
|  | TANH | 0.894±0.004 | 0.003±0.009 | 0.000±0.000 | 0.665±0.004 | 0.017±0.019 | 0.000±0.000 |
|  | SOFTPLUS | 0.899±0.002 | 0.237±0.058 | 0.016±0.021 | 0.671±0.005 | 0.261±0.042 | 0.011±0.016 |
|  | LINEAR | 0.871±0.004 | 0.113±0.066 | 0.003±0.009 | 0.653±0.005 | 0.130±0.041 | 0.003±0.006 |
|  | ELU | 0.892±0.012 | 0.113±0.082 | 0.000±0.000 | 0.672±0.004 | 0.191±0.046 | 0.006±0.010 |
| CNN-2 | RELU | 0.857±0.005 | 0.216±0.043 | 0.000±0.000 | 0.569±0.006 | 0.263±0.033 | 0.022±0.016 |
|  | EXP | 0.898±0.002 | 0.956±0.035 | 0.938±0.037 | 0.608±0.004 | 0.756±0.030 | 0.500±0.028 |
|  | SIGMOID | 0.824±0.095 | 0.206±0.078 | 0.000±0.000 | 0.571±0.003 | 0.191±0.042 | 0.000±0.000 |
|  | TANH | 0.854±0.003 | 0.184±0.066 | 0.003±0.009 | 0.576±0.008 | 0.270±0.036 | 0.003±0.006 |
|  | SOFTPLUS | 0.859±0.007 | 0.366±0.119 | 0.050±0.025 | 0.577±0.004 | 0.339±0.049 | 0.042±0.019 |
|  | LINEAR | 0.852±0.004 | 0.175±0.067 | 0.006±0.013 | 0.571±0.004 | 0.144±0.042 | 0.002±0.005 |
|  | ELU | 0.856±0.003 | 0.225±0.072 | 0.003±0.009 | 0.577±0.003 | 0.181±0.050 | 0.011±0.012 |
| CNN-50 | RELU | 0.849±0.012 | 0.900±0.064 | 0.800±0.085 | 0.566±0.012 | 0.700±0.047 | 0.378±0.059 |
|  | EXP | 0.889±0.002 | 0.825±0.035 | 0.812±0.042 | 0.616±0.005 | 0.727±0.055 | 0.495±0.039 |
|  | SIGMOID | 0.706±0.091 | 0.175±0.125 | 0.084±0.105 | 0.406±0.043 | 0.433±0.079 | 0.258±0.060 |
|  | TANH | 0.504±0.099 | 0.325±0.126 | 0.291±0.117 | 0.359±0.026 | 0.402±0.071 | 0.295±0.045 |
|  | SOFTPLUS | 0.861±0.016 | 0.881±0.056 | 0.756±0.081 | 0.585±0.007 | 0.748±0.027 | 0.481±0.039 |
|  | LINEAR | 0.846±0.013 | 0.906±0.048 | 0.791±0.054 | 0.564±0.010 | 0.680±0.050 | 0.383±0.059 |
|  | ELU | 0.847±0.012 | 0.884±0.050 | 0.809±0.081 | 0.570±0.009 | 0.688±0.064 | 0.380±0.077 |
| CNN-DEEP | MODIFIED-EXP | 0.898±0.001 | 0.284±0.051 | 0.019±0.021 | 0.663±0.006 | 0.272±0.047 | 0.019±0.022 |
|  | MODIFIED-TANH | 0.854±0.055 | 0.828±0.070 | 0.803±0.083 | 0.672±0.003 | 0.775±0.046 | 0.534±0.034 |
|  | MODIFIED-RELU | 0.900±0.002 | 0.934±0.029 | 0.912±0.027 | 0.665±0.006 | 0.836±0.041 | 0.541±0.024 |
|  | MODIFIED-SIGMOID | 0.900±0.019 | 0.934±0.038 | 0.925±0.045 | 0.668±0.004 | 0.831±0.041 | 0.569±0.035 |
|  | SUPER-RELU | 0.897±0.002 | 0.091±0.071 | 0.003±0.009 | NA | NA | NA |
| CNN-2 | MODIFIED-EXP | 0.854±0.004 | 0.306±0.071 | 0.031±0.020 | 0.598±0.002 | 0.333±0.038 | 0.037±0.017 |
|  | MODIFIED-TANH | 0.833±0.037 | 0.884±0.049 | 0.863±0.053 | 0.572±0.003 | 0.745±0.057 | 0.498±0.035 |
|  | MODIFIED-RELU | 0.881±0.015 | 0.944±0.023 | 0.919±0.021 | 0.611±0.006 | 0.752±0.037 | 0.472±0.052 |
|  | MODIFIED-SIGMOID | 0.891±0.016 | 0.953±0.032 | 0.919±0.042 | 0.608±0.004 | 0.744±0.051 | 0.498±0.040 |
|  | SUPER-RELU | 0.853±0.005 | 0.106±0.077 | 0.000±0.000 | NA | NA | NA |
| CNN-50 | MODIFIED-EXP | 0.870±0.007 | 0.887±0.049 | 0.784±0.035 | 0.606±0.008 | 0.731±0.055 | 0.425±0.031 |
|  | MODIFIED-TANH | 0.878±0.020 | 0.819±0.056 | 0.800±0.045 | 0.573±0.013 | 0.689±0.045 | 0.472±0.042 |
|  | MODIFIED-RELU | 0.878±0.009 | 0.869±0.061 | 0.831±0.069 | 0.619±0.007 | 0.775±0.032 | 0.528±0.035 |
|  | MODIFIED-SIGMOID | 0.886±0.009 | 0.863±0.029 | 0.838±0.044 | 0.616±0.006 | 0.727±0.038 | 0.492±0.043 |
|  | SUPER-RELU | 0.840±0.018 | 0.884±0.056 | 0.756±0.072 | NA | NA | NA |

To quantify how well representations learned in first layer filters match ground truth motifs embedded in the synthetic sequences, we visualized first layer filters as a position probability matrix using activation-based alignments^1, 10^ (see Methods). We employed Tomtom^34^, a motif comparison search tool, to quantify the fraction of first layer filters that yielded a statistically significant match to ground truth motifs. The filters in CNN-50, which is designed to learn whole motif representations with relu activations, were also able to capture ground truth motifs with other activation functions, with the exceptions being sigmoid and tanh activations (which is expected given their poor classification performance). CNN-2 and CNN-deep, which were designed to learn distributed representations, were unable to recover statistically significant matches to ground truth motifs for most activation functions, except the exponential activation. Notably, 36.6% of the filters of CNN-2 with softplus activations have a statistically significant match to some motif in the JASPAR database, even though the vast majority of these are not relevant for Task 1. This highlights a potential pitfall of overinterpreting filters that match a known motif in a motif database using Tomtom. Exponential activations, on the other hand, yield a match fraction of 0.953 0.038 and 0.938 0.037 to ground truth motifs for CNN-deep and CNN-2, respectively (errors represent standard deviation of the mean across 10 independent trials using different random initializations). Indeed, a qualitative comparison shows that CNN-deep’s filters visually capture many ground truth motifs when employing exponential activations (Figure 1b, see Supplemental Figs. 3 and 4 for other CNNs). Together, this demonstrates that exponential activations provide interpretable filters for CNNs, irrespective of max-pooling size and network depth without sacrificing performance.

### Generalization to real DNA sequences

To test whether exponential activations improve the interpretability of motif representations in CNN filters for real DNA sequences, we performed similar experiments on a modified DeepSea dataset^2^, truncated to include only sequences that have a peak called for at least one of 12 ChIP-seq experiments, each of which correspond to a TF in Task 1. This truncated-DeepSea dataset is similar to the synthetic dataset, except that the input sequences now have a size of 1,000 nucleotide (nt) in contrast to the 200 nt synthetic sequences in Task 1. To account for the increased complexity of features in real DNA sequences^35^, we increased the number of parameters in each hidden layer by a factor of 2. We trained each augmented CNN on the truncated-DeepSea dataset following the same protocol as Task1. Henceforth, we refer to this analysis as Task 2.

The classification performance of each CNN across different activation functions for Task 2 follow similar trends as Task 1, albeit with a larger gap between CNN-deep and the shallower CNNs (Table 2). Similarly, a comparison of the first layer filter representations demonstrate that employing exponential activations consistently leads to more interpretable filters both visually (Supplemental Figs. 5) and quantitatively (Fig. 1d) for all CNNs. For instance, CNN-deep’s filters with an exponential activation yield a match fraction of 0.859±0.038 to any JASPAR motif and a match fraction of 0.584±0.030 to a JASPAR motif associated with the ChIP-seq TFs. By contrast, CNN-deep with relu activations only yields a match fraction of 0.259±0.029 and 0.011±0.016, respectively. In general, the large decrease in motif matches to relevant motifs, *i.e.* motifs in the JASPAR database that are associated with each TF, can be attributed to the increased complexity of patterns and dependencies of motifs in real DNA sequences – here we only include positive matches to motifs that are in the JASPAR database and labelled for the ground truth TFs. Upon visual inspection, we highlight other filters of CNN-deep with exponential activations that don’t match a “relevant” motif are dedicated to other proteins, including GATA1, CTCF, GATA1-TAL1, ATF4, among many others (Supplemental Fig. 5), which were consistently found across all 10 models trained from different random initializations (see Supplemental Data).

### Transforming activations to appear exponential locally leads to interpretable motifs

To understand the properties of the exponential activation that drive improved motif representations, we transformed sigmoid, tanh, and relu activations to emulate the exponential function locally within input values in the range of −4 to 4 (Supplemental Fig. 6). This modification consists of a combination of a shift-transformation and a scale-transformation (Supplemental Table 1). Indeed, these modified activations now yield comparable motif match performance as the exponential activation for both synthetic sequences in Task 1 (Fig. 2a) and real DNA sequences in Task 2 (Table 2, Supplemental Fig. 7). As a control, we modified the exponential activation to appear relu-like (Supplemental Table 1). As expected, motif representations for CNN-2 and CNN-deep significantly decreased with the exception being CNN-50, which is designed to learn motif representations (Figs. 2a). This demonstrates that the local properties of the exponential activation can provide a strong inductive bias towards learning motif representations in first layer filters.

**Figure 2.**
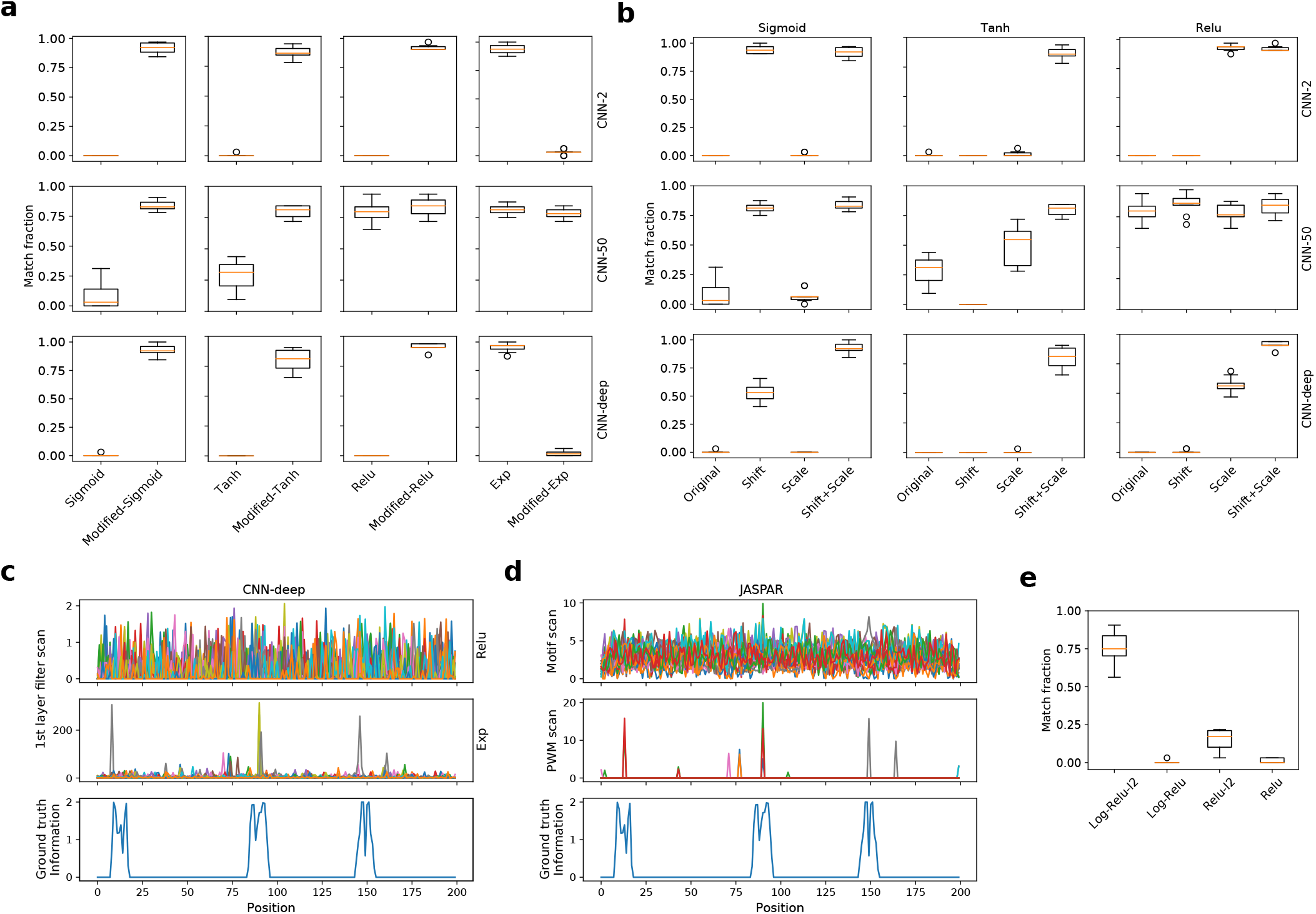
Task 1 motif representations for CNNs with modified activations. (**a**) Boxplot of the fraction of filters that match ground truth motifs for different CNNs with traditional and modified activations. (**b**) Boxplot of the fraction of filters that match ground truth motifs for an ablation study of transformations for modified activations. (**c**) First layer filter scans from CNN-deep with relu activations (top) and exponential activations (middle). Each color represents a different filter. (**d**) Motif scans (top) and PWM scans (middle) using ground truth motifs and their reverse-complements (each color represents a different filter scan). Negative PWM scan values were rectified to a value of zero. (**c**, **d**) The information content of the sequence model used to generate the synthetic sequence (ground truth), which has 3 embedded motifs centered at positions 20, 50, and 155, is shown at the bottom. (**e**) Boxplot of the fraction of filters that match ground truth motifs for CNN-deep with various activations: log activations trained with and without L2-regularization (Log-Relu-L2 and Log-Relu, respectively) and relu activations with and without L2-regularization. (**a**, **b**, **e**) Each boxplot represents the performance across 10 models trained with different random intializations (box represents first and third quartile and the red line represents the median).

Since each modified activation can be decomposed to a shift- and a scale-transformation, we employed an ablation study using CNN-deep as the base model to identify the impact of each transformation. We found that there is no consistency across transformations for different activation functions (Fig. 2b). For instance, the shift-transformation plays a stronger role for sigmoid activations, while a scale-transformation is more impactful for relu activations. Tanh activations require both scale- and shift-transformations. Here, the only constant is that stronger motif representations are consistently learned across all modified activation functions when both shift- and scale-transformations are applied.

Previously, a rectified-polynomial activation, which is inspired from dense associative memories in neuroscience^36^, was introduced in a shallow multi-layer perceptron as a mechanism to encourage hidden neurons to learn strong representations of numbers when applied to the MNIST dataset, which consists of images of handwritten digits^37^. This activation function is similar to the modified-relu, but with just a scale-transformation and a higher-order polynomial. While a rectified-polynomial activation indeed improves the motif interpretability of the shallow CNN-2 model, the effect size is significantly reduced in CNN-deep (Fig. 2b). Here, we extend the rectified-polynomial with a shift transformation, which further improves the robustness of motif representations in first layer filters across CNN architectures.

To further test whether the divergence properties of the activation function plays a larger role, we modified CNN-deep with relu activation to have a steep, linear slope of 400, we call super-relu (Supplemental Table 1). In contrast to the scale-transformation, which is a third-order polynomial, a CNN with super-relu activations do not lead to motif representations, with performance metrics that closely resembles standard relu activations (Supplemental Table 1). We also introduced and varied a scaling factor in the exponential function, *i.e.* exp(*αx*), where *α* is a scaling parameter. We found a that the scaling factor can take on a narrow range [0.5, 2] where performance for classification and motif learning in the first layer are reliable (Supplemental Table 2 and Supplemental Fig. 8). Taken together, this suggests that the divergence of the activation function alone is not a primary property that provides a strong inductive bias to learn motif representations. We surmise that the key property may be a shift from the origin combined with a sharp non-linearity that provides sensitivity to suppress background and scale up signal.

### Exponential activations are robust to initialization

Although the high divergence of the exponential function may introduce a high sensitivity to initialization, we found CNN-deep with exponential activations is robust to standard initialization strategies^38–41^ as well as over a large range of random normal initializations with varying degrees of the standard deviation (Supplemental Table 3, Supplemental Fig. 9). A large decrease in the fraction of filters that match ground truth motifs was observed as the standard deviation increases beyond 0.75, which is much larger than 0.05, the standard deviation set by He initializations for 200 nt DNA sequences^39^. Interestingly, the classification performance degraded to a much lesser extent (Supplemental Table 4).

### Scanning exponentially activated filters localizes motifs

Since our intuition suggests that the exponential activation serves to suppress background and propagate signal, the first layer filter scans may be a useful tool to reveal motif instances along a sequence, similar to PWM scans^42^. Indeed, the first layer filter scans of CNN-deep with exponential activations yield crisp peaks at locations along the DNA sequence where motifs were implanted (Fig. 2c), albeit with slight shifts that arise due to the position of the motif within each filter. By contrast, it is difficult to isolate motif instances using filters from CNN-deep with relu activations. For comparison, figure 2d shows motif scans using the ground truth motifs from the JASPAR database, given as a position probability matrix. The motif scans are very noisy, but there do seem to be some peaks with a low signal-to-noise ratio. The gold standard for motif scans are PWMs^43, 44^, which is a log-ratio of motif similarity given by the position probability matrix to background nucleotide levels. A significant improvement in sharpening the signal from the noise is evident by using the PWM transformed ground truth motif (Fig. 2d, middle row).

Since we have ground truth of the implanted motifs, the performance of locating motifs along a given sequence with motif scans can be quantified by segmenting the sequence into regions that have the implanted motif or do not and comparing the max scanned values within each region. The separation of these distributions can be summarized with the AUROC. We find that first layer filter scans of CNN-deep with exponential activations yield an AUROC of 0.889±0.201, while it is 0.391±0.331 for CNN-deep with relu activations (errors represent the standard deviation of the mean across all test sequences). Similarly, the AUROC for a motif scan represented as a position probability matrix and a PWM are 0.667 0.331 and 0.884 0.252, respectively. Here we confirm that in ideal circumstances, *i.e.* knowing the ground truth motif and background frequencies, PWM scans are a powerful approach to footprint motifs along a sequence. In practice, motifs may be nuanced from celltype to celltype^45^, requiring inference of relevant motifs specifically for a given biological system. Moreover, PWM performance is sensitive to the choice of background frequencies^46^. The appeal of CNNs is their ability to infer motif patterns and make predictions in an end-to-end fashion.

We surmise that the exponential as an activation function provides a high sensitivity which the network can exploit to suppress background and propagate signal. Similarly, a log_2_ function of the log-ratio in PWMs provides the necessary sensitivity to scale down background while maintaining signal. Both approaches serve to improve the signal-to-noise ratio. This suggests that log activations may also improve the interpretability of motif representations in first layer filters. To test this, we employed a natural log activation to first layer filters of CNN-deep. However, we found this model was not trainable – gradient descent could not minimize the loss function. This may be due to the fact that large negative values arising from noise can lead to similar issues we anticipate would occur if noise were propagated in a CNN with exponential activations. To remedy this, we applied a relu activation after the log-activation, which we call log-relu. CNN-deep with log-relu activations yielded a high classification performance (AUPR: 0.973±0.004), yet the filter representations did not recover identifiable motif representations (Fig. 2e). However, by incorporating a strong L2-regularization penalty of 0.2, we found the filters now learn improved motif representations (Fig. 2e), while maintaining a high classification performance (AUPR: 0.980±0.003). L2-regularization places a penalty on filters that have non-zero parameters. Hence, parameters are encouraged to keep background noise levels close to zero while maintaining the signals that minimize the loss, outweighing their L2 penalties. As a control, the same L2-regularization applied to CNN-deep with relu activations does not achieve a similar effect size (Fig. 2e) but still maintains a similar classification performance (AUPR: 0.978±0.003).

To further investigate whether exponential activations lead to the suppression of background and the propagation of signal, we compared pre- and post-activations for a CNN-deep model from random initialization (prior to training) and after training (Supplemental Fig. 6). At the beginning of training, the pre-activation values have a similar distribution for CNN-deep with relu and exponential activations – normally distributed with a zero mean and a standard deviation around 1.5. The post-activation values thus translate to half of the first layer neurons being activated for CNN-deep with relu activations, while they are largely in an off state (*i.e.* low values) for CNN-deep with exponential activations (Supplemental Fig. 6b). After training, the distribution of first layer neurons post-activation for CNN-deep with relu activations is decreased (Supplemental Fig. 6a), indicating that during training, the network is turning off many neurons. Interestingly, the neuron activity is largely the same before and after training for CNN-deep with relu activations. On the other hand, the distribution of first layer neurons post-activation for CNN-deep with exponential activations maintains a zero mean, but the variance increases, resulting in values that reach orders of magnitude higher up to ~400 (Supplemental Fig. 6a). This suggests that CNN-deep with exponential activations initially starts with first layer neurons effectively turned off; during training, parameters are tuned to activate just a few neurons to large values while background is maintained to low values (Fig. 2c). By contrast, CNN-deep with relu activations are initialized with a high degree of noise and so the network has to direct parameters to down-weight noise and up-weight signal. In deep CNNs, deeper layers can synergistically correct for noisy representations in the earlier layers, not requiring first layer filters to learn strong motif representations^25^.

### CNNs that learn robust motif representations are more interpretable with attribution methods

Although filter visualization is a powerful approach to assess learned representations from a CNN, they do not specify how decisions are made. Attribution methods aim to resolve this by identifying input features that are important for model predictions. To understand the role that the activation function plays in the efficacy of recovering biologically meaningful representations with attribution methods, we trained two CNNs, namely CNN-local and CNN-dist, on a synthetic regulatory classification task that serves to emulate the billboard model for cis-regulation^47, 48^. Specifically, the goal of this task, which we refer to as Task 3, is to predict whether a DNA sequence contains at least 3 “core” motifs, which are comprised of JASPAR motifs for CEBPB, Gabpa, MAX, SP1, and YY1^32^. Positive class sequences were synthesized by embedding 3 to 5 “core motifs” – randomly selected with replacement from a pool of forward- and reverse-complement core motifs – along a random sequence model. Negative class sequences were generated following the same steps with the exception that the pool of motifs include 100 non-overlapping “background motifs” from the JASPAR database^32^. Background sequences can thus contain core motifs; however, it is unlikely to randomly draw the combinations of motifs that resemble a positive regulatory code. CNN-local is a shallow network with 2 hidden layers and designed to learn interpretable filter representations with relu activations^25^, while CNN-dist is a deep network with 5 hidden layers that learn distributed representations of features. The details of the model architecture and training procedure can be found in Methods.

#### Better accuracy does not imply better interpretability

Classification performance as measured by the area under the receiver operating characteristic curve (AUC) is comparable between CNN-dist and CNN-local, with a slight edge in performance favoring CNN-dist (Fig. 3a). Since we have ground truth of which motifs were embedded and their locations in each sequence, we can test the efficacy of attribution methods. Specifically, we segment each sequence according to the information content of the sequence model into two classes – positions that are random have zero information (negative class) and positions with an embedded motif, which have information greater than zero (positive class). For each sequence, we compare the distribution of attribution scores for each class using the AUROC and AUPR to summarize how well the model correctly attributes positions in a sequence. Henceforth, we refer to this performance metric as the interpretability AUROC and AUPR.

**Figure 3.**
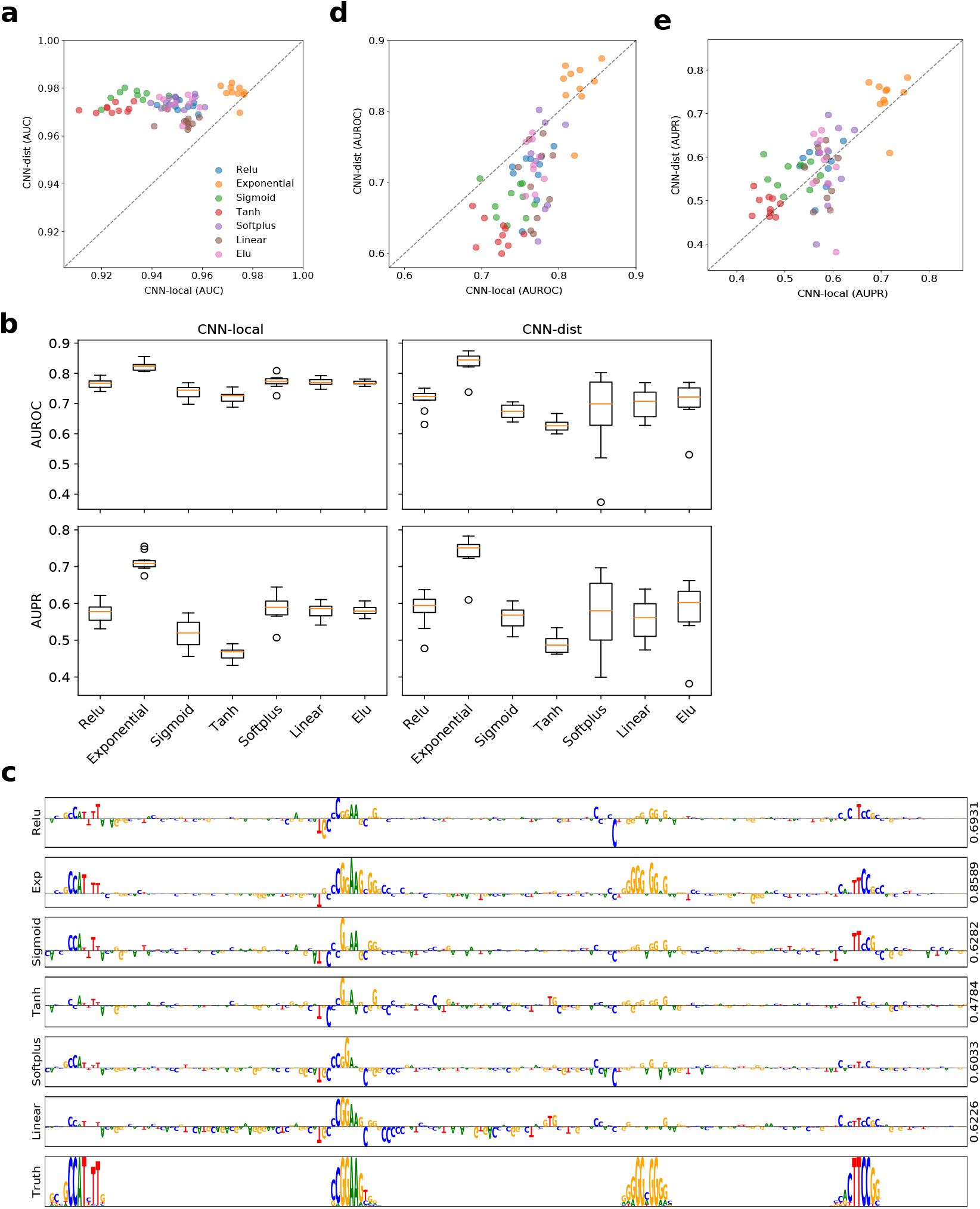
Interpretability performance of saliency maps. (**a**) Scatter plot of the classification AUC for CNN-dist versus CNN-local for different first layer activations (shown in a different color), when trained with 10 different random intializations. (**b**) Boxplot of the interpretability AUROC (top) and AUPR (bottom) for CNN-local (left) and CNN-dist (right) for different first layer activations. (**c**) Sequence logo of a saliency map for a representative test sequence generated with CNN-deep with different first layer activations (*y*-axis label). The right *y*-axis label shows the interpretability AUROC score. The sequence logo for the ground truth sequence model is shown at the bottom. Scatter plot of the (**d**) interpretability AUROC and (**e**) interpretability AUPR of saliency maps of test sequences generated from CNN-dist versus CNN-local for different activations (shown in a different color). (**a**, **d**-**e**) Each boxplot represents the performance across 10 models trained with different random intializations (box represents first and third quartile and the red line represents the median).

According to these interpretability metrics, we find that exponential activations consistently lead to the best performance for CNN-local and CNN-dist both quantitatively (Fig. 3b) and qualitatively (Fig. 3c) for attribution scores given by saliency maps, *i.e.* gradients of predictions with respect to inputs. Interestingly, CNN-local yields a slightly higher interpretability AUROC compared to CNN-dist, across most activations functions, with the exception of the exponential activation (Fig. 3d). The performance of CNN-dist yields a slightly better performance under the AUPR metric (Fig. 3e). Each metric describes slightly different aspects of interpretability. AUROC captures the ability of the network to correctly predict the embedded motifs, while penalizing spurious noise. Hence, CNN-local is less susceptible to attributing positions that are not associated with the ground truth motifs (lower false positives). AUPR considers false negative rates, and so the improved performance here suggests that CNN-dist is slightly better at capturing more ground truth patterns, while CNN-local is slightly more conservative, missing some ground truth patterns. One limitation of this study is that we are only accounting for ground truth motifs that are intentionally implanted in randomized sequence, not the spurious motifs that arise by chance (which also contributes to label noise).

The fact that CNN-dist with sigmoid activations yields high classification performance relative to CNN-local, but then yields significantly lower interpretability under both interpretability metrics, suggests that predictive performance does not necessarily imply interpretability with attribution methods. This is surprising because our intuition suggests that improved predictive models should be capturing better feature representations to explain the improved performance. The discrepancy between accurate predictions and model interpretability has also been observed in computer vision^49^.

#### Modified activations

As expected, modifying activations, such as sigmoid, tanh, and relu, which all yield low interpretability performance, to appear exponential locally significantly improves the CNN’s interpretability performance (Table 3, Supplemental Figs. 6a). Similarily, modifying the exponential to appear relu-like locally leads to a significant decrease in interpretability performance (Supplemental Fig. 7a). Moreover, we previously found that log-relu activations with an L2-regularization learn interpretable motif representations and we would expect that this activation would also lead to improved interpretability with attribution methods. Indeed, we verify this is true (Supplemental Fig. 7b). Together, this suggests that the extent that CNN filters learn robust motif representations may be indicative of network’s interpretability performance with saliency maps.

**Table 3.** Interpretability performance on Task 3. This table shows the average area under the ROC curve (AUC) classification performance, and the average AUROC and AUPR interpretability performance for CNN-dist and CNN-local for various activation functions. The errors represent the standard deviation of the mean across 10 independent trials.

| MODEL | ACTIVATION | CLASSIFICATION<br>AUC | INTERPRETABILITY |  |
| --- | --- | --- | --- | --- |
|  |  |  | AUROC | AUPR |
| CNN-DIST | RELU | 0.973±0.002 | 0.7130±0.0335 | 0.5824±0.0442 |
|  | EXP | 0.978±0.003 | 0.8352±0.0364 | 0.7362±0.0462 |
|  | SIGMOID | 0.976±0.003 | 0.6740±0.0218 | 0.5613±0.0297 |
|  | TANH | 0.971±0.002 | 0.6279±0.0195 | 0.4885±0.0226 |
|  | SOFTPLUS | 0.974±0.002 | 0.6678±0.1284 | 0.5507±0.1289 |
|  | LINEAR | 0.966±0.002 | 0.7006±0.0498 | 0.5561±0.0565 |
|  | ELU | 0.951±0.005 | 0.7065±0.0657 | 0.5814±0.0778 |
| CNN-LOCAL | RELU | 0.951±0.005 | 0.7644±0.0161 | 0.5736±0.0269 |
|  | EXP | 0.973±0.003 | 0.8245±0.0156 | 0.7125±0.0225 |
|  | SIGMOID | 0.933±0.008 | 0.7388±0.0208 | 0.5161±0.0382 |
|  | TANH | 0.924±0.006 | 0.7207±0.0193 | 0.4633±0.0183 |
|  | SOFTPLUS | 0.950±0.006 | 0.7722±0.0204 | 0.5855±0.0351 |
|  | LINEAR | 0.953±0.005 | 0.7713±0.0128 | 0.5797±0.0196 |
|  | ELU | 0.973±0.003 | 0.7686±0.0078 | 0.5813±0.0159 |
| CNN-DIST | MODIFIED-EXP | 0.973±0.003 | 0.7454±0.0253 | 0.6159±0.0317 |
|  | MODIFIED-TANH | 0.974±0.003 | 0.8128±0.0203 | 0.6829±0.0224 |
|  | MODIFIED-RELU | 0.973±0.004 | 0.7902±0.0566 | 0.6662±0.0756 |
|  | MODIFIED-SIGMOID | 0.979±0.004 | 0.8392±0.0210 | 0.7372±0.0324 |
| CNN-LOCAL | MODIFIED-EXP | 0.952±0.009 | 0.7765±0.0189 | 0.5990±0.0377 |
|  | MODIFIED-TANH | 0.972±0.005 | 0.8150±0.0188 | 0.6900±0.0386 |
|  | MODIFIED-RELU | 0.970±0.002 | 0.8236±0.0121 | 0.7107±0.0157 |
|  | MODIFIED-SIGMOID | 0.972±0.002 | 0.8214±0.0117 | 0.7100±0.0179 |

#### Other attribution methods

Many issues have been documented for saliency maps^8, 22–24^, and hence the poor performance may be a reflection of a flawed methodology and not necessarily the interpretability of the model. By comparing the interpretability performance of different attribution methods, including *in silico* mutagenesis, integrated gradients, and DeepSHAP, we find that different attribution methods yield a very different ability to recover ground truth motifs (Fig. 4, a-b and Supplemental Table 5). The gold standard is *in silico* mutagenesis, and it consistently yields the most reliable attribution maps, with DeepSHAP in second place. Irrespective of the attribution method, CNNs that employ exponential activations significantly improve performance consistently across all interpretability metrics compared to other activations, both quantitatively (Fig. 4, a-b) and qualitatively (Fig. 4, c-d).

**Figure 4.**
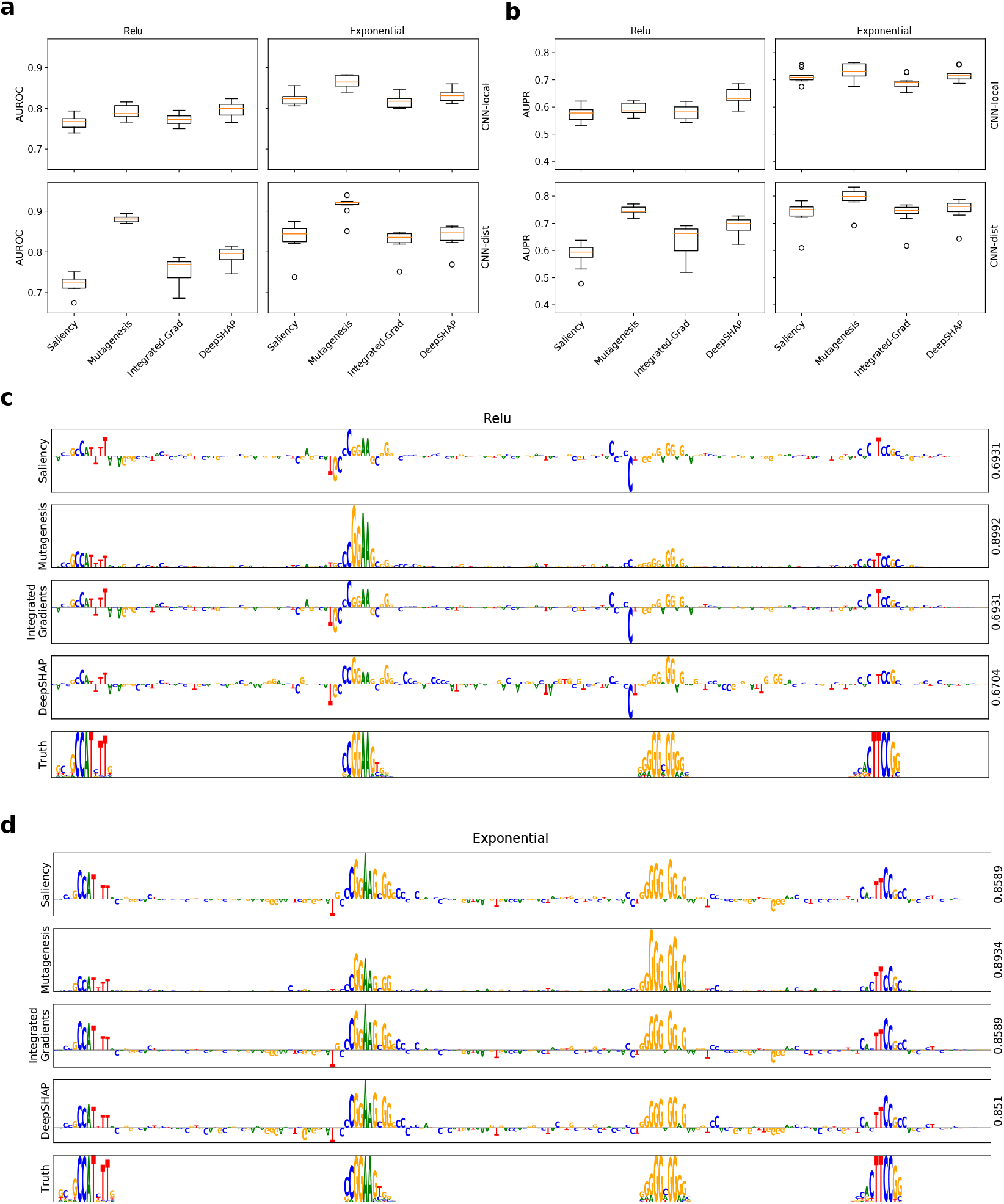
Interpretability performance comparison of different attribution methods. Boxplots of the interpretability AUROC (a) and AUPR (**b**) for CNN-local (top) and CNN-dist (bottom) with relu activations (left) and exponential activations (right) for different attribution methods. Each boxplot represents the performance across 10 models trained with different random intializations (box represents first and third quartile and the red line represents the median). Sequence logo for a saliency map for a Task 3 test sequence generated with different attribution methods for CNN-deep with relu activations (**c**) and exponential activations (**d**). The right *y*-axis label shows the interpretability AUROC score. (**c**-**d**) The sequence logo for the ground truth sequence model is shown at the bottom.

### Attribution methods on *in vivo* regulatory genomic sequences

To demonstrate whether exponential activations improve the quality of representations from attribution methods on real regulatory genomic sequences, we trained two Basset models on a multi-task classification of chromatin accessibility sites from the Basset dataset^1^, which we refer to as Task 4. Each Basset model consists of 3 convolutional layers followed by 2 fully-connected hidden layers (see Methods for details), with the only difference being the first layer activations, relu or exponential. Similarly, both Basset models yield very similar classification performance with an AUPR of 0.486±0.042 and 0.489±0.041 for relu and exponential activations, respectively. As expected, figure 5a anecdotally shows that Basset with exponential activations leads to more interpretable motif representations for an accessible site in fibroblast cells, uncovering cleaner representations of 3 motifs – TCF4, NFIX, and HLF – which are all import regulators previously identified for chromatin accessibility^50–52^. In general, a qualitative comparison shows that a Basset model with exponential activations leads to less noisy representations with saliency maps (Supplemental Fig. 8). While we cannot assess the importance of the spurious attribution scores here, recall that CNNs with relu activations tend to yield spurious attribution scores on synthetic data, which do not correspond to ground truth motif patterns. This is consistent with the behavior that we observe in real DNA sequences. Moreover, a comparison of the first layer filter representations to the JASPAR database shows that exponential activations result in a match fraction of 0.617, while relu activations yielded 0.370. Visual inspection of the filter representations showed that Basset with exponential activations contains more information content with better visual matches to known motifs in the JASPAR database (see Supplemental data).

**Figure 5.**
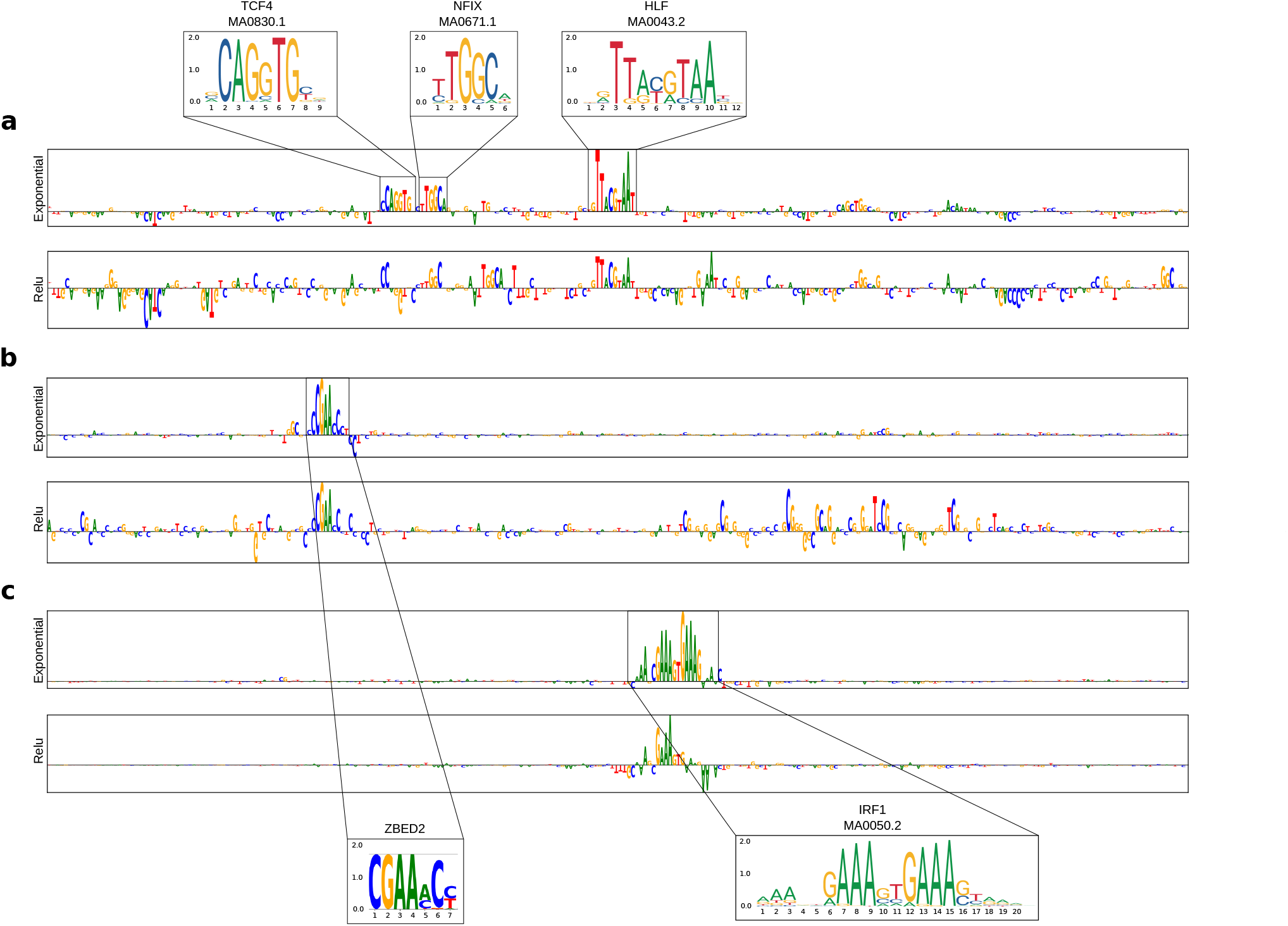
Attribution score comparison for real regulatory DNA sequences. Sequence logo of a saliency map generated for a representative test sequence from a CNN with exponential activations (top) and relu activations (bottom) trained on (**a**) Task 4 (DNase-seq peaks), (**b**) Task 5 (ZBED2 ChIP-seq peaks) and (**c**) Task 6 (IRF1 ChIP-seq peaks). The sequence logo of the known motifs are highlighted.

We also employed ResidualBind, a CNN originally employed for RNA-protein interactions^53^, to classify positive-label DNA sequences about ChIP-seq peaks for ZBED2 (Task 5) and IRF1 (Task 6) in pancreatic ductal adenocarcinoma cells versus negative-label sequences about ChIP-seq peaks for H3K27ac marks within the same cell-type (nonoverlapping peaks with the TFs)^54^. In each task, we trained two nearly identical ResidualBind models, with the only difference being the first layer activation was either relu or exponential. Figure 5b-c, shows that ResidualBind with relu activations result in seemingly noisier attribution scores with spurious nucleotides given importance scores, compared to ResidualBind with exponential activations (see Supplemental Figs. 9 and 10 for more examples). Classification performance was comparable across all CNNs for each task. For Task 5, ResidualBind yielded an AUROC of 0.882 and 0.898 with relu and exponential activations, respectively; for Task 6, the AUROC was 0.985 and 0.982.

## Discussion

A major draw of deep learning in genomics is their powerful ability to automatically learn features from the data that enable it to make accurate predictions. In biology, it is critical that we understand what features it has learned in order to build trust in these black box predictive models and to potentially gain new biological insights from them. Model interpretability is key to understanding these features. Deep CNNs, however, tend to learn distributed representations of sequence motifs that are not necessarily human interpretable. Although attribution methods aim to identify features that affect decision making, in practice, their scores tend to be noisy and difficult to interpret. Here, we show that an exponential activation applied to the first layer is a powerful approach to encourage first layer filters to learn sequence motifs and also to improve the efficacy of attribution scores, revealing more human interpretable representations.

One major consequence from this study raises the red flag that a CNN that yields high classification performance does not necessarily provide meaningful representations with attribution methods. Previous studies have focused on comparing representations from different attribution methods using only a single model^8^. Here, we show that different models, each with comparable classification performance, can yield significantly different representations with the same attribution methods. We believe that deeper CNNs may be learning a noisier representation of biological signals but still capable of making equally good (if not better) predictions on held-out test data. Investigating properties of the underlying function that enable good generalization performance but poor interpretability is important to address the root of this issue.

### Other activation functions

While we showed that local properties of the exponential activation lead to interpretable motifs in first layer filters, it remains unclear to what extent each property matters – level of divergence and the shift from the origin. Moreover, other alternatives to the exponential have not been explored here. A simple alternative of the exponential is the power series expansion of polynomials, which can improve computational efficiency. Another major factor that was not explored here is the initialization strategy, which can have a great influence on the trainability and performance of deep CNNs^33^.

### Limitations to filter interpretability

Although one of the goals of this paper is to design CNNs such that their filters are more interpretable, there are many issues that must be considered so as not to over-interpret filter representations. A filter’s information content is not a useful property for its importance as is the usage of the filter. Low information content filters will undoubtedly have higher usage than high-information content filters, and how they are weighted, and hence their importance, is contained in deeper layers. Filter influence is a promising direction to uncover the effect size that a filter has on model predictions, by performing an intervention that effectively turns off all significant activations^55^. However, this too can be challenging to fully understand because of the complex dependencies between filters in deeper layers.

Moreover, designing CNNs with interpretable filters sets a hard limit on the number and sizes of patterns that can be detected. Assembling motifs hierarchically through partial motifs exponentially increases the expressive power to build more representations, thus reducing the sensitivity to the numbers and sizes of the filters. However, we showed a strong association that attribution methods are noisier when partial motifs are learned, perhaps a result of assembling noisier representations of motifs in deeper layers.

Attribution methods provide a single-nucleotide resolution map of the importance of each nucleotide in a given sequence, which often highlights representations that resemble known motifs, confirming our belief that the network has learned meaningful biology. However, attribution methods explored here are first-order interpretability methods, defining the importance of individual nucleotides – not the importance of the entire motif on model predictions. Although recent progress is extending these class of methods to second-order attributions^53, 56–58^, they cannot uncover the effect size of motifs on model predictions. Global interpretability analysis via *in silico* experiments is one avenue that shows great promise in uncovering the importance of whole features^59^.

### Scoring mutations

Scoring the functional consequence of mutations is one promising application of deep learning in genomics. However, in order to move forward, we must trust that the deep learning model is making reliable predictions. Testing model predictions on held-out test data is not sufficient to evaluate whether we can trust model predictions for single nucleotide mutations. We demonstrated that models that yield high classification performance can yield very low interpretability with first-order attribution methods, including *in silico* mutagenesis. While we do not elucidate all of the factors that underlie this discrepancy between model classification and interpretability, we identified a strong association that models that learn more robust representations of motifs in first layer filters lead to significantly improved interpretability with first-order attribution methods. We suspect that if a robust motif representation is learned anywhere in the network, then attribution methods will perform well. Hence, verifying that a network has learned a strong motif representations can serve as a necessary (but not sufficient) quality control to ensure trust in attribution methods, including *in silico* mutagenesis. Since the representations of filters in deeper layers of a CNN are challenging to recover, enforcing that first layer filters learn strong motif representations can be achieved and easily verified with exponential (or equivalent) activations.

### Tradeoff no more

Interpretability of first layer convolutional filters is seemingly at odds with classification performance, especially for deeper networks, which are more flexible in terms of the function classes that they can fit^27^. Previously, we showed one design principle for CNNs to learn interpretable motif representations is to increase the max-pool size after the first convolutional layer^25^. This of course comes at the sacrifice of network depth, which is generally associated with better classification performance. Here, we show that CNNs with exponential activations substantially improve motif representations in the first layer while not making any sacrifices in performance. Importantly, this trick can be applied to networks of any depth. Although not tested here, we believe that it could also improve filter interpretability in deeper layers to potentially capture motif-motif interactions. In practice, the exponential should probably only be applied to one layer for numerical stability. One possible solution is to instead employ an exponential equivalent that doesn’t diverge, such as a modified-sigmoid activation, which is bounded, making it possible to explore “interpretable” activations in multiple layers.

### Identifiability

Interpretability of model parameters is challenging because of a lack of identifiability in over-parameterized models – the representations in filters result in a different set of representations when trained from different initializations. While exponential activations do not completely solve this problem, we find that the similarity in representations learned by first layer filters tend to improve models trained with different random intializations, hence making their parameters more robust and identifiable.

### Robustness and interpretability

Small perturbations to the data, such as single nucleotide mutations, can have a dramatic impact on model predictions. In computer vision, targeted adversarial perturbations can lead to egregious classification errors^60^. The fragility of attribution methods has also been demonstrated with adversarial examples^61, 62^. Adversarial training has emerged to remedy the fragility of CNN predictions^63^. Unexpectedly, adversarial training leads to some intriguing properties, such as a smoother fitted function^49^ and improved interepretability with attribution methods^64^. Early work has also demonstrated the benefit of adversarial training to improve the interpretability of attribution methods in genomics^65^. Recently, in computer vision, it has been shown that adversarial training promotes learning more robust features in contrast to low-rank noise features that are correlated with each class and intrinsic to datasets^66^. It remains unclear whether the contrapositive statement is true: if a network learns robust features, then it will be adversarially robust. Here, we empirically demonstrate the relationship between CNNs learning robust features (via design principles) and improved interpretability (which is a property that follows from adversarial robustness). The causal relationship between robust features, interpretability, adversarial robustness remains a topic for further investigation.

## Methods

### Data

#### Task 1

Task 1 consists of a multi-task classification dataset from Ref.^25^. This dataset consists of 30,000 synthetic DNA sequences embedded with known transcription factor motifs. Synthetic sequences, each 200 nucleotides long, were sampled from a uniform (*i.e.* equiprobable) sequence model implanted with 1 to 5 known TF motifs, randomly selected with replacement from a pool of 12 motifs, which include Arid3, CEBPB, FOSL1, Gabpa, MAFK, MAX, MEF2A, NFYB, SP1, SRF, STAT1, and YY1. Sequences were sampled once from a unique sequence model. This dataset makes a simplifying assumption that the only important pattern for a given binding event is the presence of a PWM-like motif in a sequence. The dataset is randomly split to a training, validation, and test set according to the fractions 0.7, 0.1, and 0.2, respectively.

#### Task 2

Task 2 consists of a truncated version of the DeepSea dataset^2^. The DeepSea dataset was reduced to 12 labels by removing sequences that did not correspond to 12 class labels defined in Supplemental Table 1 in Ref.^25^. This truncation only includes 12 labels that match the TFs in Task 1 in K562 cells. Sequences are 1000 nucleotides in length.

#### Task 3

We generated 20,000 synthetic sequences each 200 nts long by embedding known motifs in specific combinations in a uniform sequence model. Positive class sequences were synthesized by sampling a sequence model embedded with 3 to 5 “core motifs” – randomly selected with replacement from a pool of 10 position frequency matrices, which include the forward and reverse-complement motifs for CEBPB, Gabpa, MAX, SP1, and YY1^32^ – along a random sequence model. Negative class sequences were generated following the same steps with the exception that the pool of motifs include 100 non-overlapping “background motifs” from the JASPAR database^32^. Background sequences can thus contain core motifs; however, it is unlikely to randomly draw motifs that resemble a positive regulatory code. We randomly combined synthetic sequences of the positive and negative class and randomly split the dataset into training, validation and test sets with a 0.7, 0.1, and 0.2 split, respectively.

#### Task 4

Task 4 sequences are from the Basset dataset^1^. This includes 164 DNase-seq datasets from ENCODE^67^ and Roadmaps Epigenomics^68^. The processed dataset consists of 1,879,982 training and 71,886 test sequences that are 600 nts long. Each sequence has an associated binary label vector corresponding to the presence of a statistically significant peak for each of the 164 cell types.

#### Tasks 5 and 6

Processed ZBED2 and IRF1 ChIP-seq data for Tasks 5 and 6 were acquired from^54^. Positive class sequences were defined as 400 nt sequences centered on ChIP-seq peaks in pancreatic ductal adenocarcinoma cells. Negative class sequences were defined as 200 nt sequences centered on peaks for H3K27ac ChIP-seq peaks that do not overlap with any positive peaks from the same cell type. We randomly subsampled the negative class sequences to balance the class labels. We randomly split the dataset into training, validation and test sets with a 0.7, 0.1, and 0.2 split, respectively. The total number of sequences is 4,902 and 3,892 for Tasks 5 and 6, respectively. We augmented the training data by generating reverse-complement sequences.

### Models

#### Task 1

CNN-2, CNN-50, and CNN-deep take as input a 1-dimensional one-hot-encoded sequence with 4 channels, one for each nucleotide (A, C, G, T), and have a fully-connected (dense) output layer with 12 neurons that use sigmoid activations. The hidden layers for each model are:

1. CNN-2
  1. convolution (32 filters, size 19, activation) max-pooling (size 2)
  2. convolution (124 filters, size 5, relu) max-pooling (size 50)
  3. fully-connected layer (512 units, relu)
2. CNN-50
  1. convolution (32 filters, size 19, activation) max-pooling (size 50)
  2. convolution (124 filters, size 5, relu) max-pooling (size 2)
  3. fully-connected layer (512 units, relu)
3. CNN-deep
  1. convolution (32 filters, size 19, activation)
  2. convolution (48 filters, size 9, relu) max-pooling (size 4)
  3. convolution (96 filters, size 6, relu) max-pooling (size 4)
  4. convolution (128 filters, size 4, relu) max-pooling (size 3)
  5. fully-connected layer (512 units, relu)

All models incorporate batch normalization^69^ in each hidden layer prior to the nonlinear activation; dropout^70^ with probabilities corresponding to 0.1 (layer 1), 0.1 (layer 2), 0.5 (layer 3) for CNN-2 and CNN-50; and 0.1 (layer 1), 0.2 (layer 2), 0.3 (layer 3), 0.4 (layer 4), 0.5 (layer 5) for CNN-deep; and *L*2-regularization on all parameters in the network with a strength equal to 1e-6, unless stated otherwise.

#### Task 2

Same models as Task 1 but with augmented hidden layers, multiplying the number of filters or hidden units by a factor of 2. Note that the inputs to the models also change from 200 nt to 1000 nt.

#### Task 3

We designed two CNNs, namely CNN-local and CNN-deep, to learn “local” representations (whole motifs) and “distributed” representations (partial motifs), respectively. Both take as input a 1-dimensional one-hot-encoded sequence (200 nt) and have a fully-connected (dense) output layer with a single sigmoid activation. The hidden layers for each model are:

1. CNN-local
  1. convolution (24 filters, size 19, activation) max-pooling (size 50)
  2. fully-connected layer (96 units, relu)
2. CNN-dist
  1. convolution (24 filters, size 7, activation)
  2. convolution (32 filters, size 9, relu) max-pooling (size 3)
  3. convolution (48 filters, size 6, relu) max-pooling (size 4)
  4. convolution (64 filters, size 4, relu) max-pooling (size 3)
  5. fully-connected layer (96 units, relu)

We incorporate batch normalization in each hidden layer prior to the nonlinear activation; dropout with probabilities corresponding to: CNN-local (layer1 0.1, layer2 0.5) and CNN-deep (layer1 0.1, layer2 0.2, layer3 0.3, layer4 0.4, layer5 0.5); and *L*2-regularization on all parameters in the network with a strength equal to 1e-6.

#### Task 6

We replicated a Basset-like model that takes as input a 1-dimensional one-hot-encoded sequence (600 nt) and have a fully-connected (dense) output layer with 164 units with sigmoid activations. The hidden layers for each model are:

1. Basset
  1. convolution (300 filters, size 19, activation) max-pooling (size 3)
  2. convolution (200 filters, size 11, relu) max-pooling (size 4)
  3. convolution (200 filters, size 7, relu) max-pooling (size 4)
  4. fully-connected (1000 units, relu)
  5. fully-connected (1000 units, relu)

We incorporate batch normalization in each hidden layer prior to the nonlinear activation; dropout with probabilities corresponding to: 0.2, 0.2, 0.2, 0.5 and 0.5; and *L*2-regularization on all parameters in the network with a strength equal to 1e-6.

#### Tasks 5 and 6

We employed a Residualbind-like model that takes as input one-hot encoded sequence (400 nt) and have a fully-connected layer to a single unit with sigmoid activations. The hidden layers are:

1. Residualbind
  1. convolution (24 filters, size 19, activation) residual block max-pooling (size 10)
  2. convolution (48 filters, size 7, relu) max-pooling (size 5)
  3. convolution (64 filters, size 7, relu) max-pooling (size 4)
  4. fully-connected (96 units, relu)

The residual block consists of a convolutional layer with filter size 5, followed by batch normalization, relu activation, dropout with a probability of 0.1, convolutional layer with filter size 5, batch normalization, and a element-wise sum with the inputs to the residual block, a so-called skipped connection, followed by a relu activation, and dropout with a probability of 0.2. For each hidden layer, we incorporate batch normalization^69^ and dropout^70^ with probabilities corresponding to: 0.1, 0.3, 0.4, and 0.5.

#### Training

We uniformly trained each model by minimizing the binary cross-entropy loss function with mini-batch stochastic gradient descent (100 sequences) for 100 epochs with Adam updates using default parameters^71^. We decayed the learning rate which started at 0.001, and when the performance metric that was monitored (AUPR for Tasks 1, 2, 4; AUROC for Tasks 3, 5, 6) did not improve for 5 epochs, the learning rate was decayed by a factor 0.3. All reported performance metrics are drawn from the test set using the model parameters which yielded the highest performance metric on the validation set. Each model was trained (10 times for Tasks 1-3 and once for Task 4-6) with different random initializations according to Ref.^38^.

### Filter analysis

#### Filter visualization

To visualize first layer filters, we scanned each filter across every sequence in the test set. Sequences whose maximum activation was less than a cutoff of 50% of the maximum possible activation achievable for that filter in the test set were removed^1, 10^. A subsequence the size of the filter centered about the max activation for each remaining sequence and assembled into an alignment. Subsequences that are shorter than the filter size due to their max activation being too close to the ends of the sequence were also discarded. A position frequency matrix was then created from the alignment and converted to a sequence logo using Logomaker^72^. The motif representations were largely not sensitive to the activation threshold (Supplemental Fig. 15), with a slight increase in motif matches to ground truth motifs for CNN-deep with relu activations trained on Task 1 and a decrease for exponential activations, which arises due to the reduced sequence diversity in the alignment for high thresholds.

#### Quantitative motif comparison

The interpretability of each filter was assessed using the Tomtom motif comparison search tool^34^ to determine statistically significant matches to the 2016 JASPAR vertebrates database^32^, with the exception of Grembl, for which many filters yielded a statistically significant match, despite visually appearing non-informative. Since the ground truth motifs are available for our synthetic dataset, we can test whether the CNNs have captured *relevant* motifs.

#### Motif localization analysis

The performance of locating motifs along a given sequence with motif scans was quantified by segmenting the sequence into regions that have the implanted motif or do not. This was determined by calculating the information content of the sequence model used to generate the synthetic sequence and segmenting ground truth from background according to an information content threshold greater than zero. A buffer size of 10 nts was added to the boundaries of each embedded motif, because motif positions within filters are not necessarily centered. The max filter scan score was given for each segmented region with a label of one for ground truth regions and a label of zero otherwise. The positive and negative label scores were aggregated across all test sequences and the AUROC was calculated.

### Attribution analysis

#### Attribution methods

To test interpretability of trained models, we generate attribution scores by employing saliency maps^6^, *in silico* mutagenesis^1, 2, 10^, integrated gradients^7^, and DeepSHAP^9^. Saliency maps were calculated by computing the gradients of the predictions with respect to the inputs. Integrated gradients were calculated by adding the saliency maps generated from 20 sequences that linearly interpolate between a reference sequence and the query sequence. We average the integrated gradients score across 10 different reference sequences generated from random shuffles of the query sequence. For DeepSHAP, we used the package from Ref.^9^, and averaged the attribution scores across 10 different randomly shuffled reference sequences. We found that 10 randomly shuffled reference sequences marked the elbow point where the inclusion of additional sequences only provided a marginal improvement in performance (Supplemental Fig. 16). Saliency maps, integrated gradients, and DeepShap scores were multiplied by the query sequence (times inputs). In silico mutagenesis was calculated by generating new sequences with all possible single nucleotide mutations of a sequence and monitoring the change in prediction compared to wildtype. *In silico* mutagenesis scores were reduced to a single score for each position by calculating the L2-norm of the mutagenesis scores across nucleotides for each position. All attribution maps were visualized as a sequence logo using Logomaker^72^.

#### Quantifying interpretability

Since we have the ground truth of embedded motif locations in each sequence, we can test the efficacy of attribution scores. To quantify the interpretability of a given attribution map, we calculate the area under the receiver-operating characteristic curve (AUROC) and the area under the precision-recall curve (AUPR), comparing the distribution of attribution scores where ground truth motifs have been implanted (positive class) and the distribution of attribution scores at positions not associated with any ground truth motifs (negative class). Specifically, we first multiply the attribution scores (*S*_*ij*_) and the input sequence (*X*_*ij*_) and reduce the dimensions to get a single score per position, according to *C*_*i*_ = ∑ _*j*_ *S*_*ij*_ *X* _*ij*_, where *j* is the alphabet and *i* is the position. We then calculate the information of the sequence model, *M* _*ij*_, according to *I* _*i*_ = log_2_ 4 - ∑ _*j*_ *M* _*ij*_ log_2_ *M*_*ij*_. Positions that are given a positive label are defined by *I*_*i*_ *>* 0, while negative labels are given by *I*_*i*_ = 0. The AUROC and AUPR is then calculated separately for each sequence using the distribution of *C*_*i*_ at positive label positions against negative label positions.

## Availability

Dataset and code: github.com/koo-lab/exponential_activations

## Acknowledgements

This work was supported in part by funding from the NCI Cancer Center Support Grant (CA045508) and the Simons Center for Quantitative Biology at Cold Spring Harbor Laboratory. MP was supported by NIH NCI RFA-CA-19-002. The authors would like to thank Dimitri Krotov, who provided inspiration for the exponential activation. We would also like to thank Justin Kinney, Antonio Majdandzic, and Ammar Tareen for helpful discussions.

## Author contributions statement

P.K.K. conceived of the experiments. P.K.K. and M.P. conducted the experiments. P.K.K. and M.P. analyzed the results. All authors reviewed the manuscript.

---

**Supplemental Table 1.** Modified activation functions. This table shows the activation functions for **z**, which represents the pre-activation values in a hidden layer.

| Activation | Function |
| --- | --- |
| Super-relu | $\max[400 * \mathbf{z}, 0]$ |
| Modified-relu | $\text{relu}((\mathbf{z} - 0.2)^3)$ |
| Modified-exp | $\max[\exp(0.001 * \mathbf{z}), 0]$ |
| Modified-sigmoid | $4000 * \text{sigmoid}(\mathbf{z} - 8)$ |
| Modified-tanh | $500 * \tanh(\mathbf{z} - 6) + 500$ |
| Log-relu | $\max[\log(\mathbf{z}), 0]$ |
| Shift-sigmoid | $\text{sigmoid}(\mathbf{z} - 8)$ |
| Scale-sigmoid | $4000 * \text{sigmoid}(\mathbf{z})$ |
| Shift-tanh | $\tanh(\mathbf{z} - 6)$ |
| Scale-tanh | $500 * \tanh(\mathbf{z}) + 500$ |
| Shift-relu | $\text{relu}((\mathbf{z} - 0.2))$ |
| Scale-relu | $\text{relu}(\mathbf{z}^3)$ |

**Supplemental Table 2.**
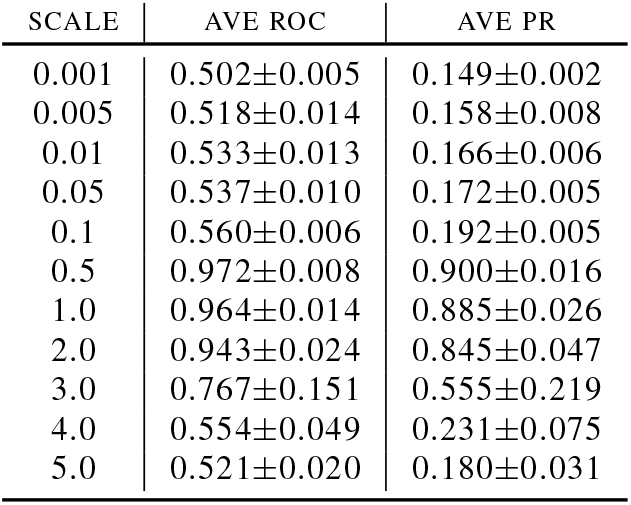
Task 1 classification performance comparison of CNN-deep with exponential activations with different scaling factors. This table shows the average area under the receiver operating characteristic curve (AUROC) and the average area under the precision recall curve (AUPR) for different exponential scaling factors for CNN-deep, *i.e.* exp(*α***z**), where *α* is the scaling factor. The errors represent the standard deviation of the mean across 10 independent trials.

**Supplemental Table 3.** Task 1 classification performance comparison of CNN-deep with different random initializations. This table shows the average area under the receiver operating characteristic curve (AUROC) and the average area under the precision recall curve (AUPR) for CNN-deep with relu and exponential activations trained with different standard initializations. The errors represent the standard deviation of the mean across 10 independent trials.

| INITIALIZATION | RELU |  | EXPONENTIAL |  |
| --- | --- | --- | --- | --- |
|  | AVE ROC | AVE PR | AVE ROC | AVE PR |
| GLOROT-NORMAL | 0.974±0.001 | 0.901±0.001 | 0.970±0.011 | 0.896±0.020 |
| GLOROT-UNIFORM | 0.974±0.001 | 0.901±0.001 | 0.973±0.008 | 0.901±0.017 |
| HE-NORMAL | 0.974±0.001 | 0.900±0.002 | 0.974±0.002 | 0.903±0.007 |
| HE-UNIFORM | 0.974±0.001 | 0.899±0.001 | 0.975±0.001 | 0.905±0.002 |
| LECUN-NORMAL | 0.973±0.002 | 0.898±0.005 | 0.972±0.008 | 0.900±0.015 |
| LECUN-UNIFORM | 0.973±0.001 | 0.899±0.002 | 0.970±0.011 | 0.896±0.019 |

**Supplemental Table 4.** Task 1 classification performance comparison of CNN-deep with random normal initialization with different standard deviations. This table shows the average area under the receiver operating characteristic curve (AUROC) and the average area under the precision recall curve (AUPR) for CNN-deep with relu and exponential activations trained with random normal initialization with different standard deviations. The errors represent the standard deviation of the mean across 10 independent trials.

| SCALE | RELU |  | EXPONENTIAL |  |
| --- | --- | --- | --- | --- |
|  | AVE ROC | AVE PR | AVE ROC | AVE PR |
| 0.001 | 0.974±0.000 | 0.901±0.002 | 0.962±0.018 | 0.880±0.035 |
| 0.005 | 0.974±0.001 | 0.901±0.002 | 0.966±0.014 | 0.886±0.032 |
| 0.01 | 0.974±0.000 | 0.902±0.001 | 0.972±0.009 | 0.899±0.017 |
| 0.05 | 0.974±0.000 | 0.900±0.001 | 0.970±0.011 | 0.896±0.020 |
| 0.1 | 0.974±0.001 | 0.900±0.002 | 0.969±0.012 | 0.894±0.023 |
| 0.2 | 0.972±0.003 | 0.896±0.007 | 0.975±0.001 | 0.906±0.001 |
| 0.3 | 0.972±0.001 | 0.897±0.002 | 0.974±0.002 | 0.903±0.004 |
| 0.4 | 0.972±0.001 | 0.897±0.002 | 0.975±0.001 | 0.904±0.002 |
| 0.5 | 0.969±0.006 | 0.890±0.012 | 0.973±0.002 | 0.901±0.004 |
| 0.75 | 0.970±0.002 | 0.891±0.006 | 0.971±0.002 | 0.895±0.006 |
| 1.0 | 0.968±0.002 | 0.889±0.002 | 0.969±0.002 | 0.889±0.004 |
| 2.0 | 0.963±0.006 | 0.877±0.015 | 0.967±0.001 | 0.884±0.003 |
| 3.0 | 0.962±0.003 | 0.875±0.007 | 0.961±0.002 | 0.871±0.004 |
| 4.0 | 0.959±0.008 | 0.868±0.017 | 0.959±0.003 | 0.867±0.007 |
| 5.0 | 0.961±0.002 | 0.871±0.004 | 0.958±0.003 | 0.864±0.007 |

**Supplemental Table 5.** Interpretability AUROC comparison of attribution methods on Task 3. This table shows the average AUROC interpretability performance (top) and the AUPR interpretability performance (bottom) for different attribution methods for CNN-dist and CNN-local for various activation functions. The errors represent the standard deviation of the mean across 10 independent trials.

|  | MODEL | ACTIVATION | SALIENCY | *In silico* MUTAGENESIS | INTEGRATED GRADIENTS | DEEPSHAP |
| --- | --- | --- | --- | --- | --- | --- |
| AUROC | CNN-DIST | RELU | 0.7130±0.0335 | 0.8803±0.0076 | 0.7542±0.0316 | 0.7904±0.0200 |
|  |  | EXP | 0.8352±0.0364 | 0.9136±0.0227 | 0.8275±0.0274 | 0.8385±0.0270 |
|  |  | SIGMOID | 0.6740±0.0218 | 0.8960±0.0063 | 0.7327±0.0159 | 0.7993±0.0150 |
|  |  | TANH | 0.6279±0.0195 | 0.8768±0.0034 | 0.6875±0.0166 | 0.7558±0.0123 |
|  |  | SOFTPLUS | 0.6678±0.1284 | 0.8916±0.0037 | 0.6996±0.1302 | 0.7674±0.0774 |
|  |  | LINEAR | 0.7006±0.0498 | 0.8596±0.0097 | 0.7530±0.0349 | 0.7822±0.0213 |
|  |  | ELU | 0.7065±0.0657 | 0.8800±0.0066 | 0.7420±0.0616 | 0.7891±0.0316 |
|  | CNN-LOCAL | RELU | 0.7644±0.0161 | 0.7914±0.0158 | 0.7713±0.0140 | 0.7962±0.0175 |
|  |  | EXP | 0.8245±0.0156 | 0.8650±0.0148 | 0.8178±0.0156 | 0.8315±0.0157 |
|  |  | SIGMOID | 0.7388±0.0208 | 0.7573±0.0356 | 0.7441±0.0166 | 0.7703±0.0208 |
|  |  | TANH | 0.7207±0.0193 | 0.7369±0.0145 | 0.7347±0.0157 | 0.7623±0.0219 |
|  |  | SOFTPLUS | 0.7722±0.0204 | 0.8029±0.0171 | 0.7784±0.0168 | 0.8004±0.0188 |
|  |  | LINEAR | 0.7713±0.0128 | 0.7961±0.0117 | 0.7742±0.0110 | 0.8010±0.0114 |
|  |  | ELU | 0.7686±0.0078 | 0.7942±0.0125 | 0.7739±0.0079 | 0.7956±0.0113 |
| AUPR | CNN-DIST | RELU | 0.5824±0.0442 | 0.7462±0.0157 | 0.6362±0.0561 | 0.6898±0.0307 |
|  |  | EXP | 0.7362±0.0462 | 0.7930±0.0377 | 0.7350±0.0417 | 0.7499±0.0394 |
|  |  | SIGMOID | 0.5613±0.0297 | 0.7703±0.0216 | 0.6291±0.0261 | 0.7203±0.0182 |
|  |  | TANH | 0.4885±0.0226 | 0.7389±0.0131 | 0.5439±0.0271 | 0.6531±0.0153 |
|  |  | SOFTPLUS | 0.5507±0.1289 | 0.7680±0.0122 | 0.5755±0.1557 | 0.6734±0.0864 |
|  |  | LINEAR | 0.5561±0.0565 | 0.7047±0.0368 | 0.6061±0.0525 | 0.6634±0.0228 |
|  |  | ELU | 0.5814±0.0778 | 0.7374±0.0281 | 0.6205±0.0862 | 0.6897±0.0415 |
|  | CNN-LOCAL | RELU | 0.5736±0.0269 | 0.5916±0.0219 | 0.5805±0.0257 | 0.6374±0.0306 |
|  |  | EXP | 0.7125±0.0225 | 0.7319±0.0274 | 0.6893±0.0239 | 0.7185±0.0218 |
|  |  | SIGMOID | 0.5161±0.0382 | 0.5335±0.0717 | 0.5172±0.0310 | 0.5734±0.0368 |
|  |  | TANH | 0.4633±0.0183 | 0.4921±0.0321 | 0.4872±0.0197 | 0.5592±0.0287 |
|  |  | SOFTPLUS | 0.5855±0.0351 | 0.6122±0.0280 | 0.5901±0.0299 | 0.6407±0.0328 |
|  |  | LINEAR | 0.5797±0.0196 | 0.5954±0.0206 | 0.5775±0.0177 | 0.6416±0.0206 |
|  |  | ELU | 0.5813±0.0159 | 0.5937±0.0232 | 0.5849±0.0176 | 0.6357±0.0227 |

**Supplemental Figure 1.**
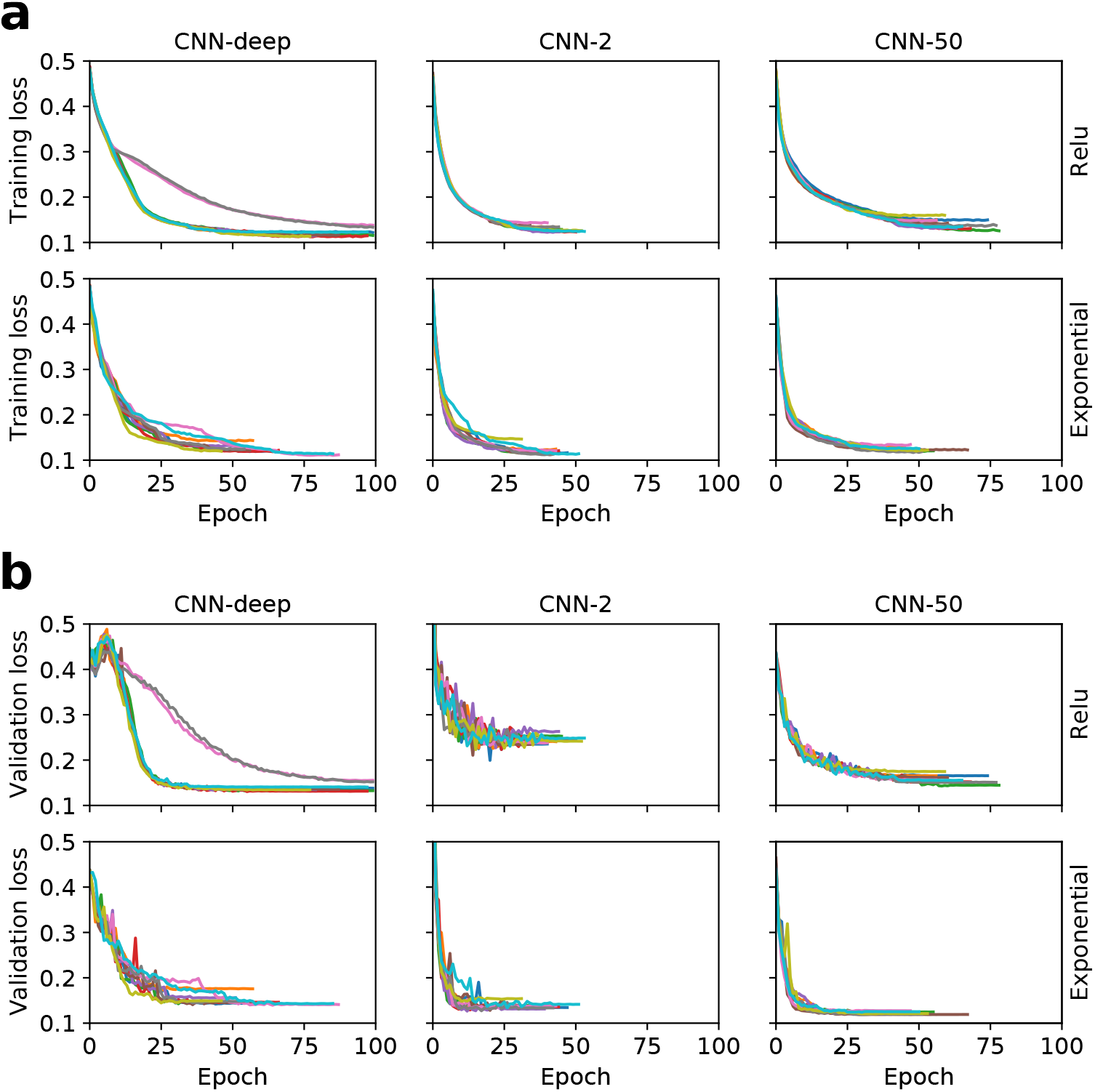
Training stability for Task 1. Loss for each epoch for CNNs with relu activations (top) and exponential activations (bottom) for (a) training data and (b) validation data. Each curve represents a different trial using different random initializations (shown in a different color).

**Supplemental Figure 2.**
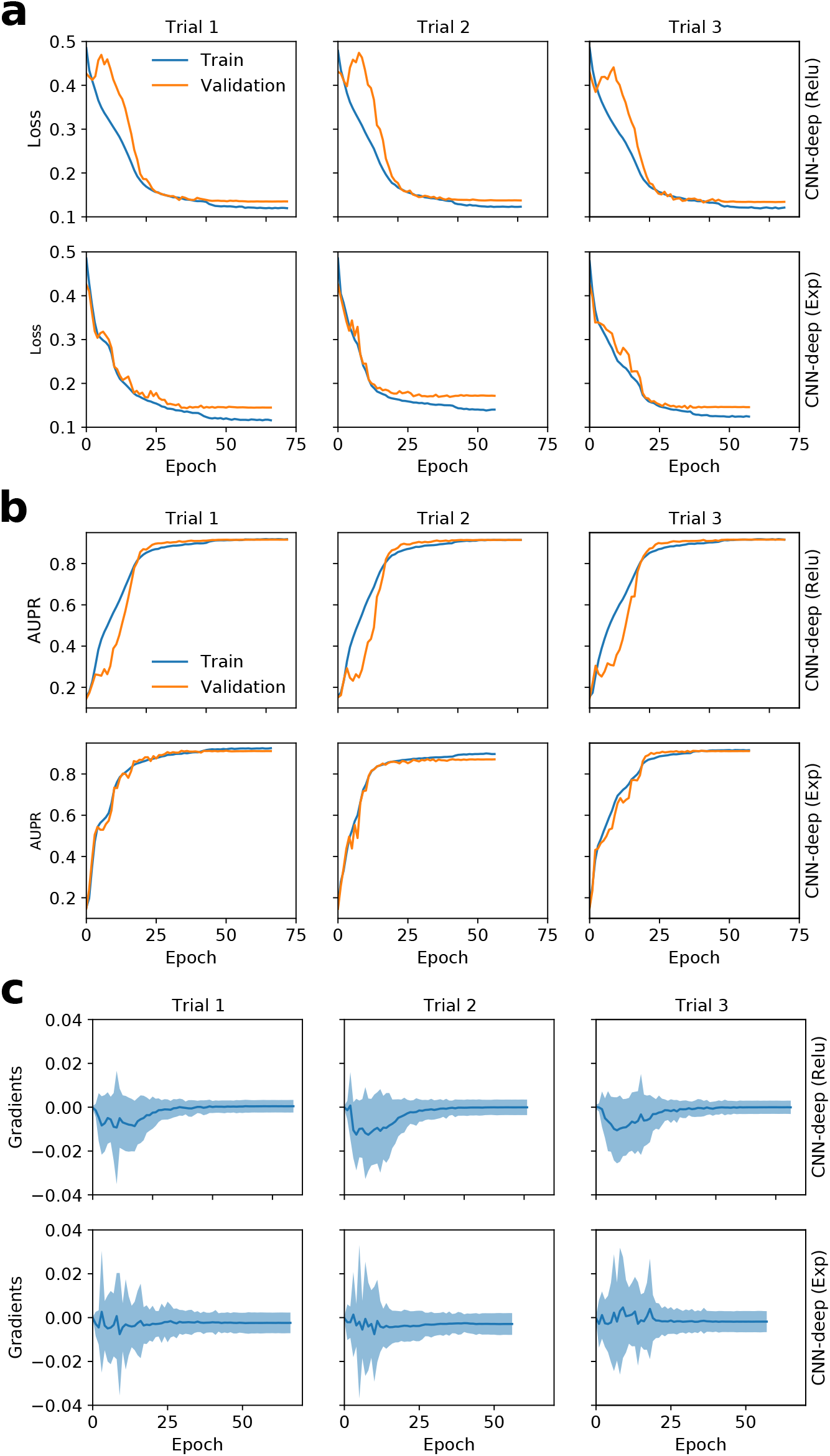
Stability of CNN-deep training on Task 1. (a) Training loss (blue) and validation loss (orange) for each epoch of training for CNN-deep with relu activations (top) and exponential activations (bottom) for different random initializations (*i.e.* different trials). Different trials stop short of 100 training steps due to early stopping. (b) The area under the precision recall curve (AUPR) and (c) the average gradients to the first layer filters are shown for the same models indicated by the trial. Shaded boundaries in (c) represent the standard deviation of the mean.

**Supplemental Figure 3.**
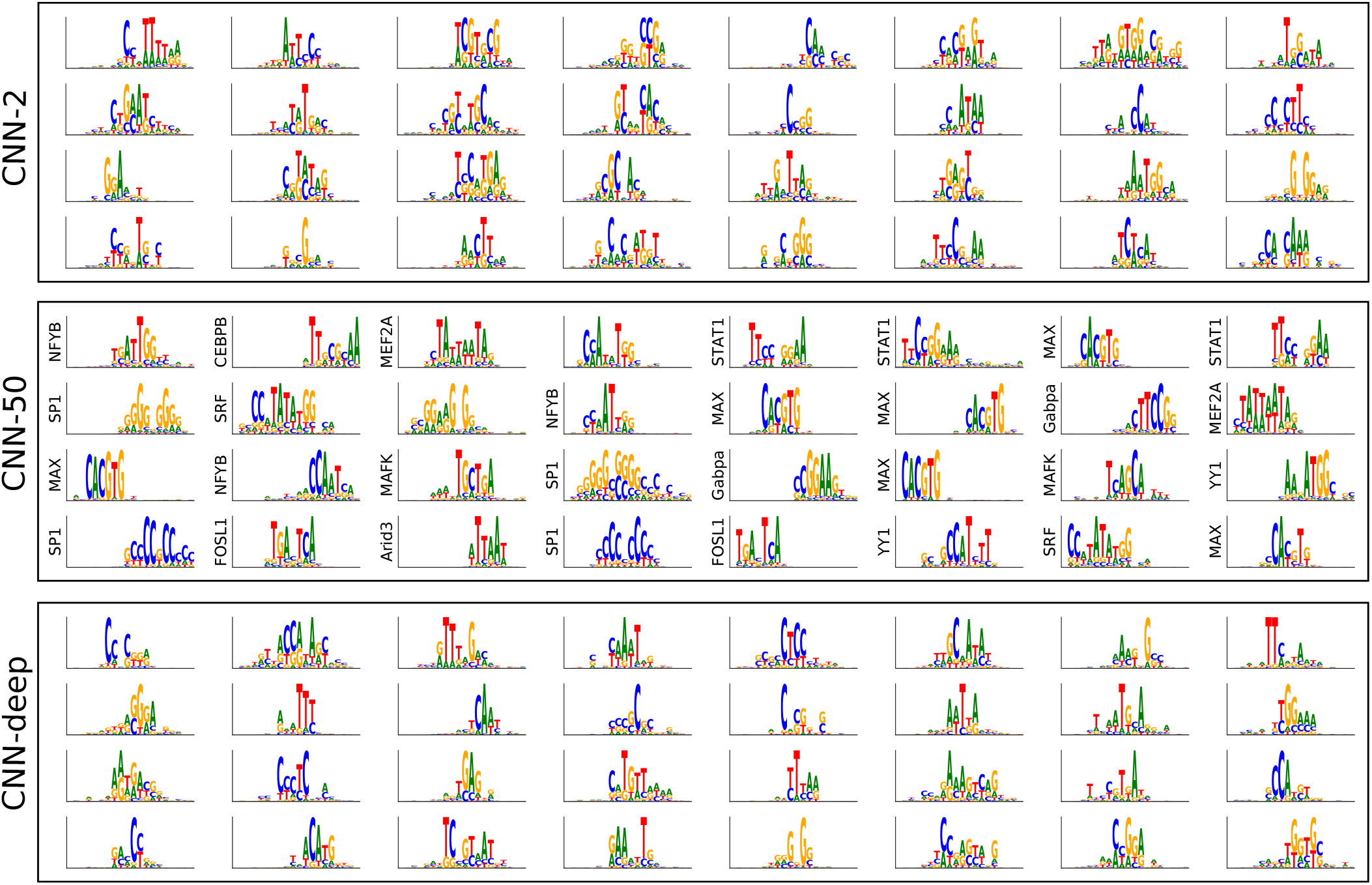
Task 1 filter comparison for CNNs with relu activations. Sequence logos for first convolutional layer filters are shown for CNN-2 (top), CNN-50 (middle) and CNN-deep (bottom) with relu activations. The *y*-axis label on select filters represents a statistically significant match to a ground truth motif as determined by Tomtom with an E-value threshold of 0.1.

**Supplemental Figure 4.**
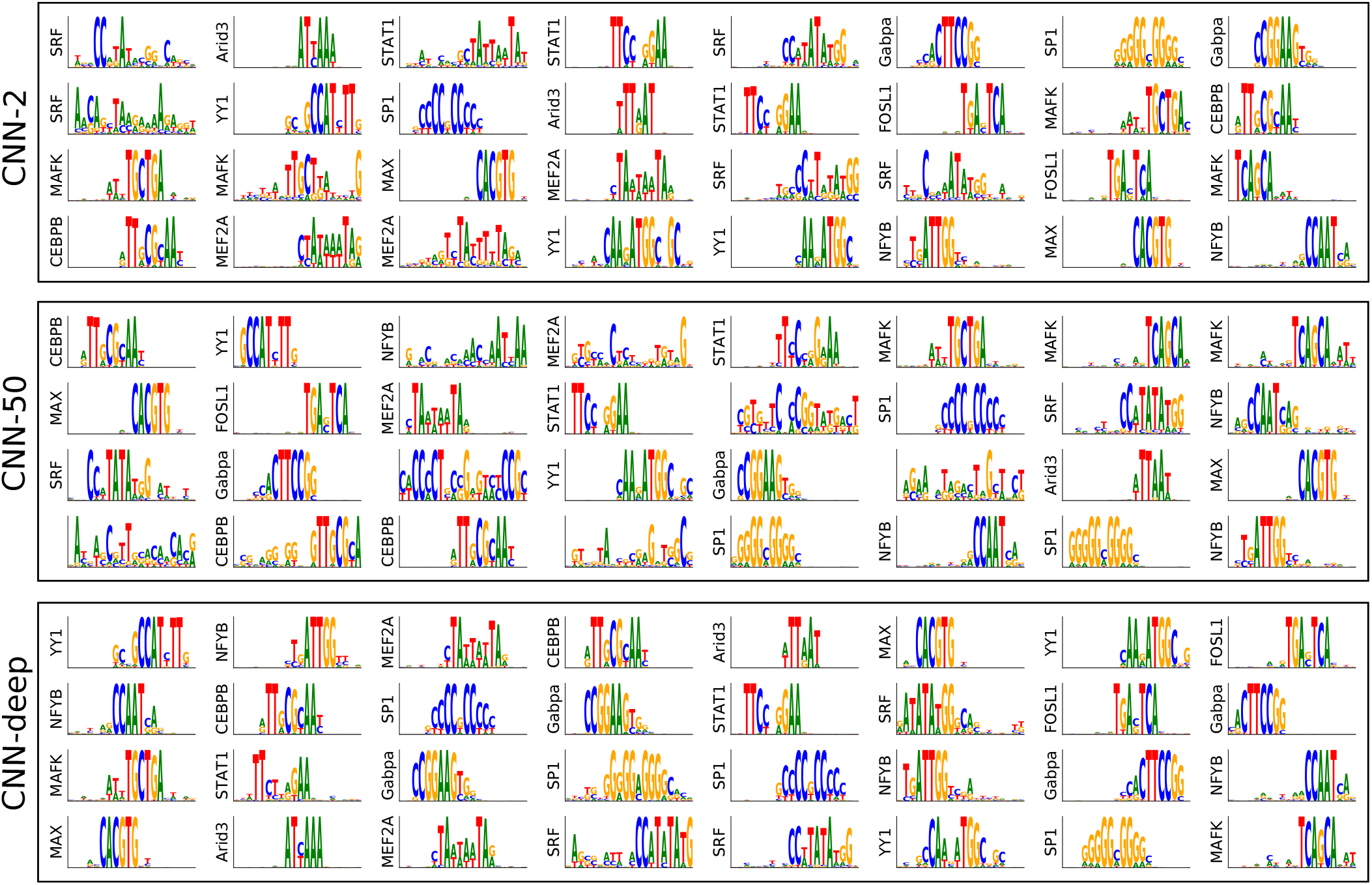
Task 1 filter comparison for CNNs with first layer exponential activations. Sequence logos for first convolutional layer filters are shown for CNN-2 (top), CNN-50 (middle) and CNN-deep (bottom) with exponential activations. The *y*-axis label on select filters represents a statistically significant match to a ground truth motif as determined by Tomtom with an E-value threshold of 0.1.

**Supplemental Figure 5.**
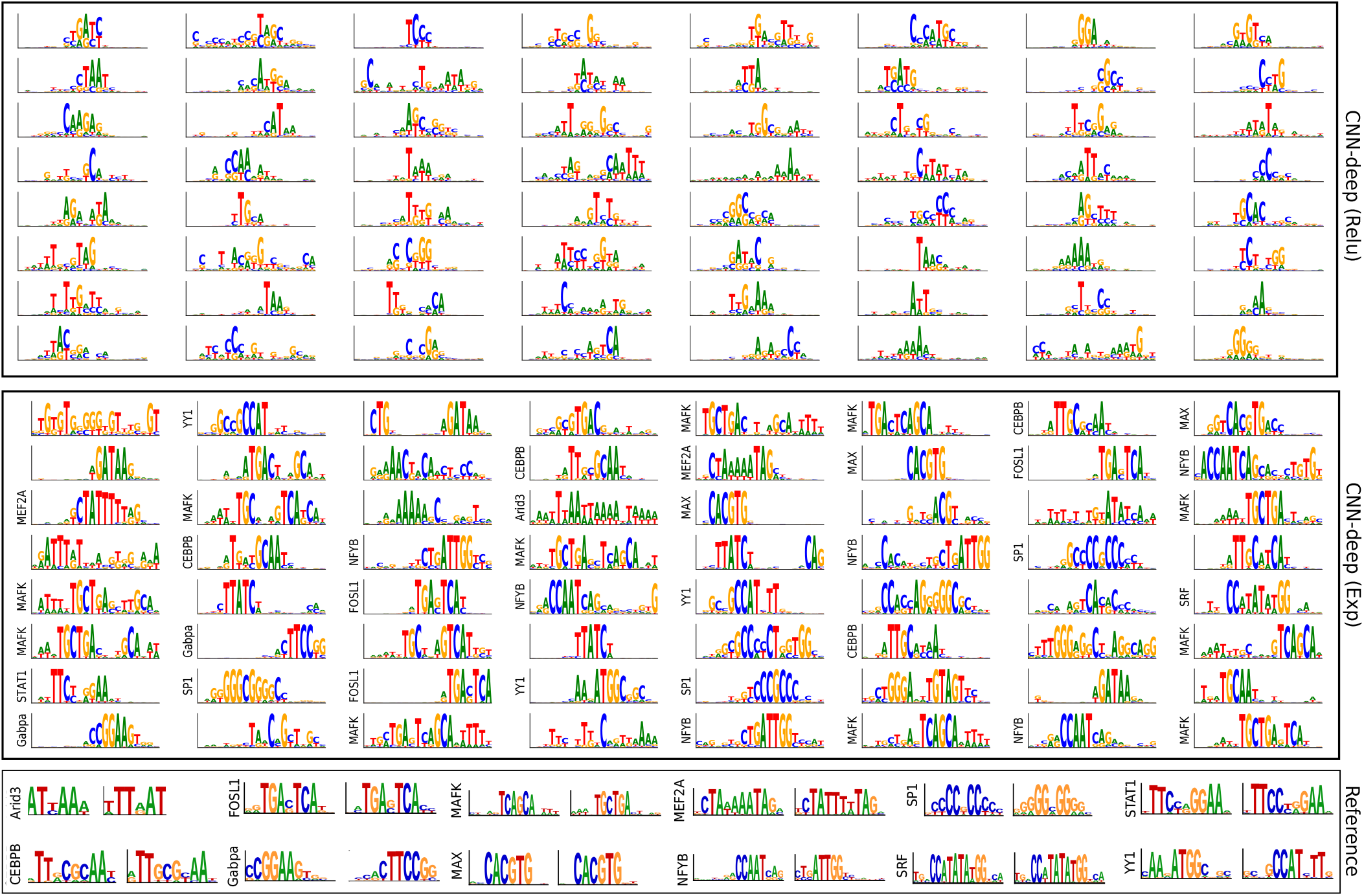
Task 2 filter comparison for relu and exponential activations. Sequence logos for first convolutional layer filters are shown for CNN-deep with relu activations (top) and exponential activations (middle). The sequence logo of the known motifs and its reverse complement for corresponding TFs in the JASPAR database are shown below. The *y*-axis label on select filters represents a statistically significant match to any reference TF motif in the JASPAR database as determined by Tomtom with an E-value threshold of 0.1.

**Supplemental Figure 6.**
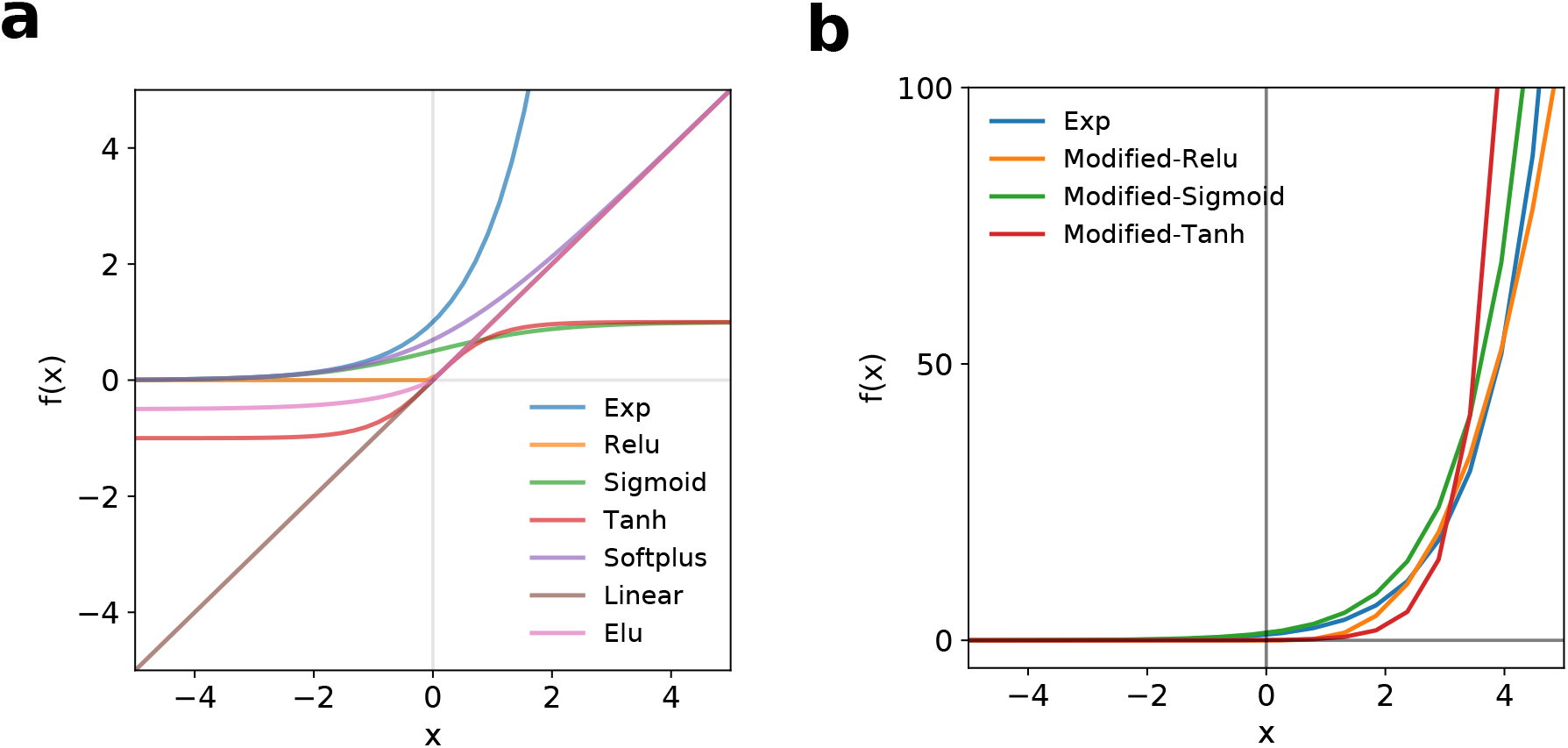
Activations. (**a**) Plot of commonly used activation functions, including exponential (exp), relu, sigmoid, tanh, softplus, linear, and elu. (**b**) Plot of modified activation functions for relu, sigmoid, and tanh, which are similar to the exponential locally for inputs in the range of −4 to 4.

**Supplemental Figure 7.**
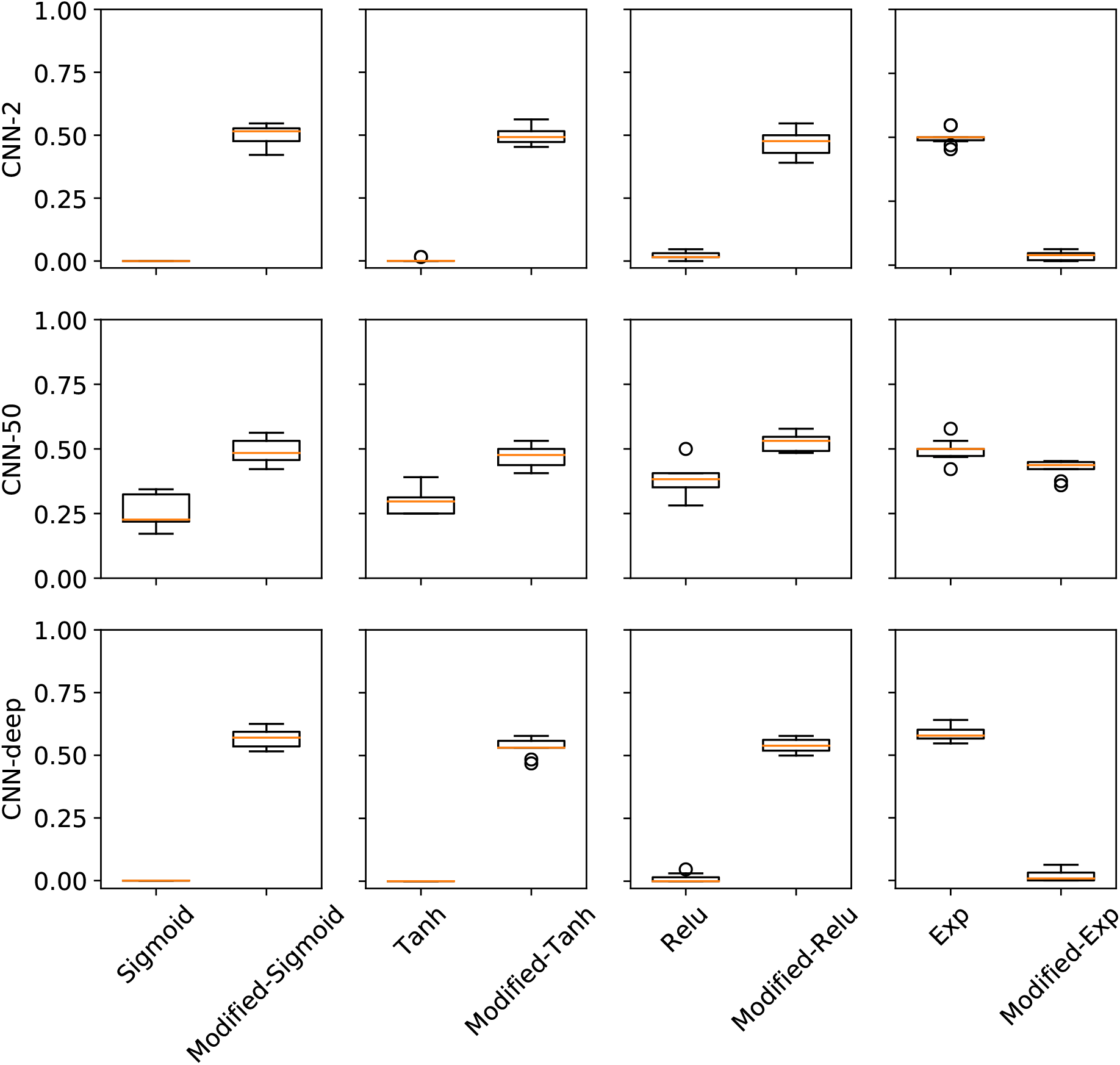
Task 2 filter performance for modified activations. Box plot of the fraction of filters that match ground truth motifs for various first layer activations and modified activations in CNN-2 (top), CNN-50 (middle), and CNN-deep (bottom) trained on real DNA sequences of Task 2. (**c**-**d**) Each box plot represents the performance across 10 models trained with different random initializations (box represents first and third quartile and the red line represents the median).

**Supplemental Figure 8.**
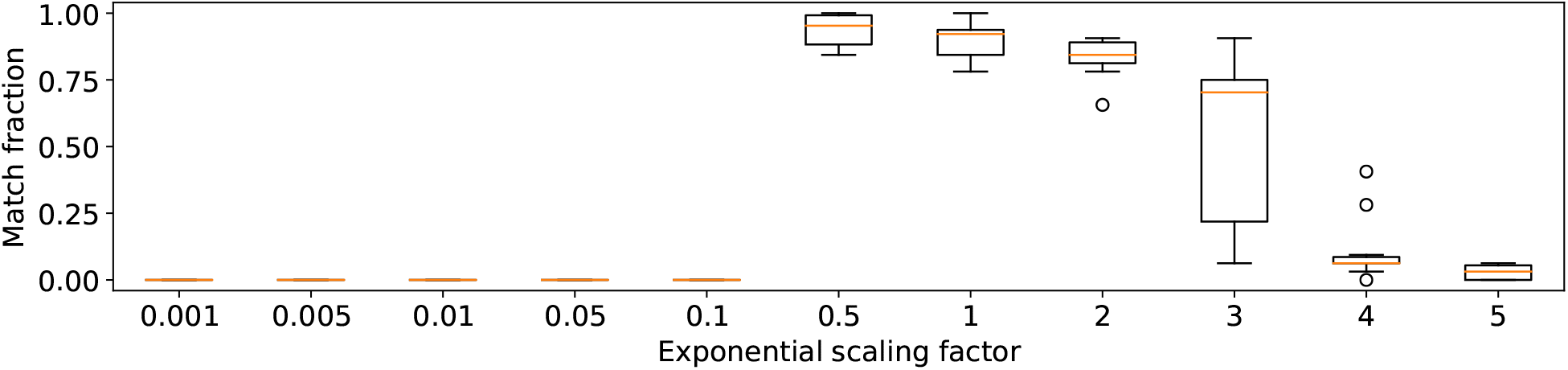
Task 1 filter performance for CNN-deep with different exponential scale factors. Box plot of the fraction of filters that match ground truth motifs for CNN-deep with different exponential scaling factors, *i.e.* exp(*α***z**), where *α* is the scaling factor. Each box plot represents the performance across 10 models trained with different random initializations (box represents first and third quartile and the red line represents the median).

**Supplemental Figure 9.**
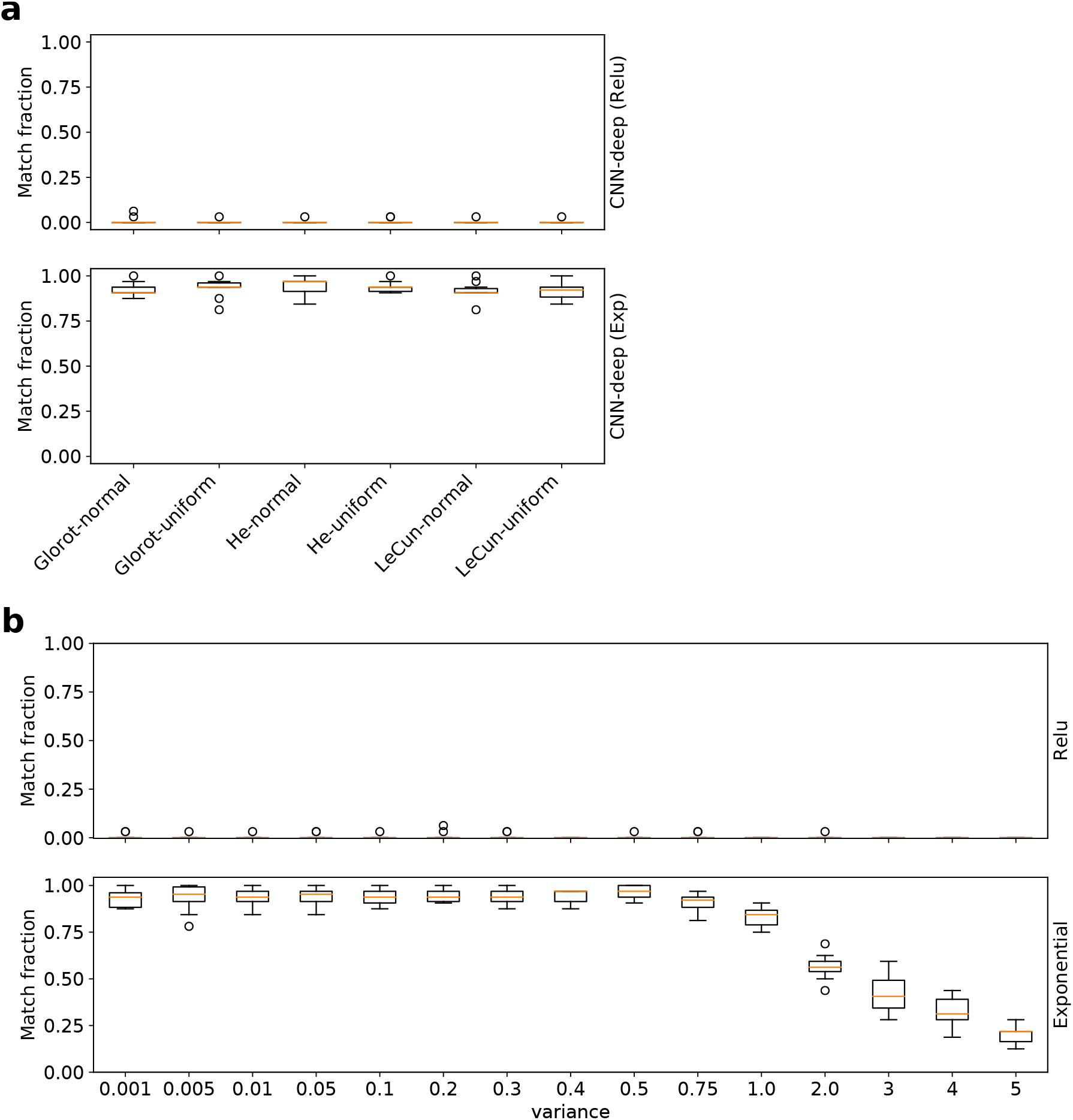
Task 1 filter performance comparison for CNN-deep with different initializations. Box plot of the fraction of filters that match ground truth motifs for CNN-deep with relu activations (top) and exponential activations (bottom) for (a) different standard random initializations and (b) random normal initializations with zero mean and varying standard deviations. Each box plot represents the performance across 10 models trained with different random initializations (box represents first and third quartile and the red line represents the median).

**Supplemental Figure 10.**
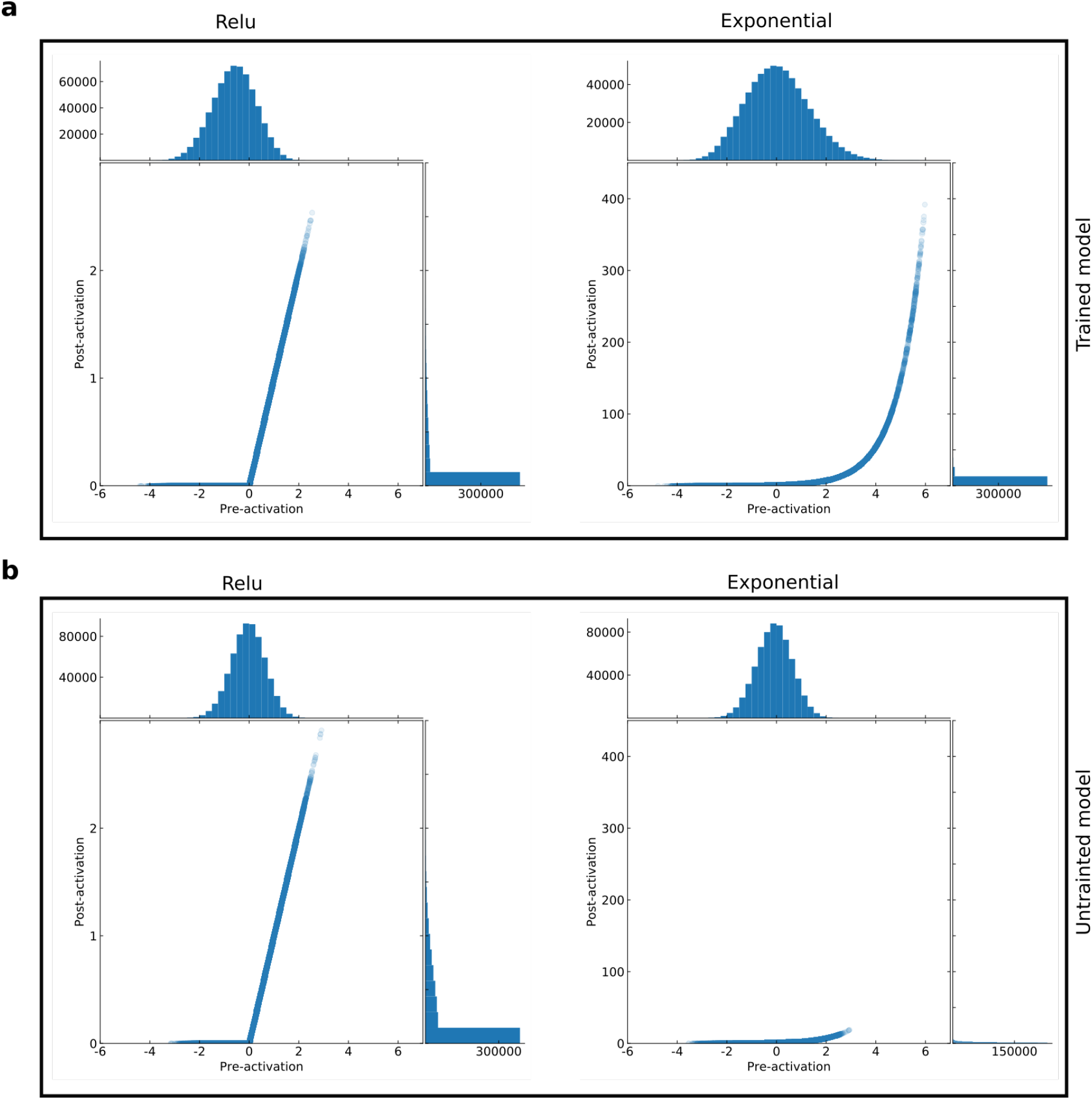
Comparison of pre- and post-activations. (**a**) Shows a scatter plot of the pre-activation values and the post-activation values for 100 random test sequences for a trained CNN-deep with relu activations (left) and exponential activations (right). The top histogram shows the pre-activation values and the histogram on the left shows the post-activation values. (**b**) shows a similar plot but with an untrained CNN-deep model – just random initialization.

**Supplemental Figure 11.**
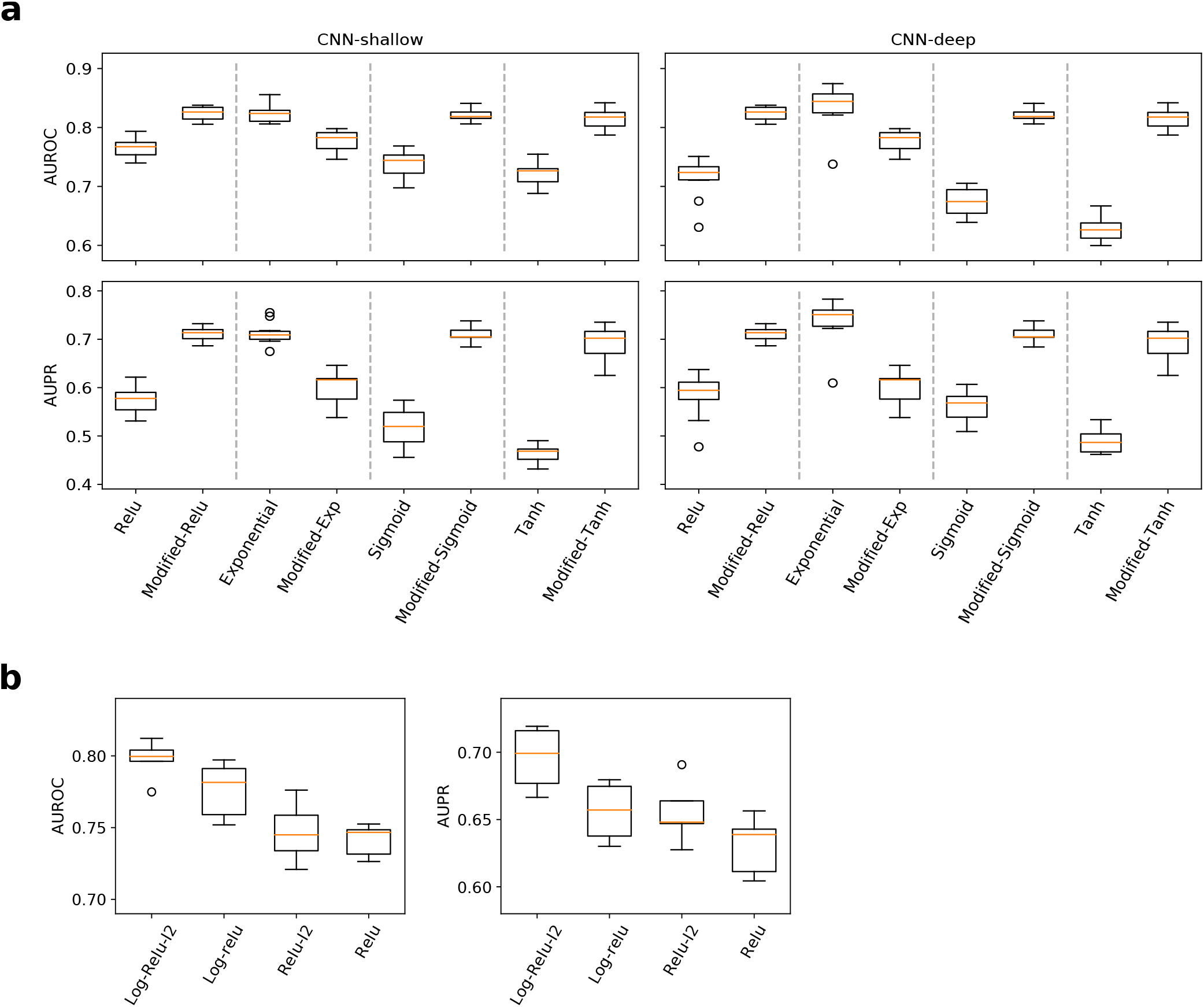
Task 3 attribution performance for CNNs with modified activations. (**a**) Box plots of the interpretability AUROC (top) and AUPR (bottom) for CNN-local (left) and CNN-dist (right) with various activations. (**b**) Box plots of the interpretability AUROC (left) and AUPR (right) for CNN-dist with log-relu activation with L2-regularization (Log-Relu-l2) and without (Log-Relu) and relu activations with L2-regularization (Relu-l2) and without (Relu). Each box plot represents the performance across 10 models trained with different random initializations (box represents first and third quartile and the red line represents the median).

**Supplemental Figure 12.**
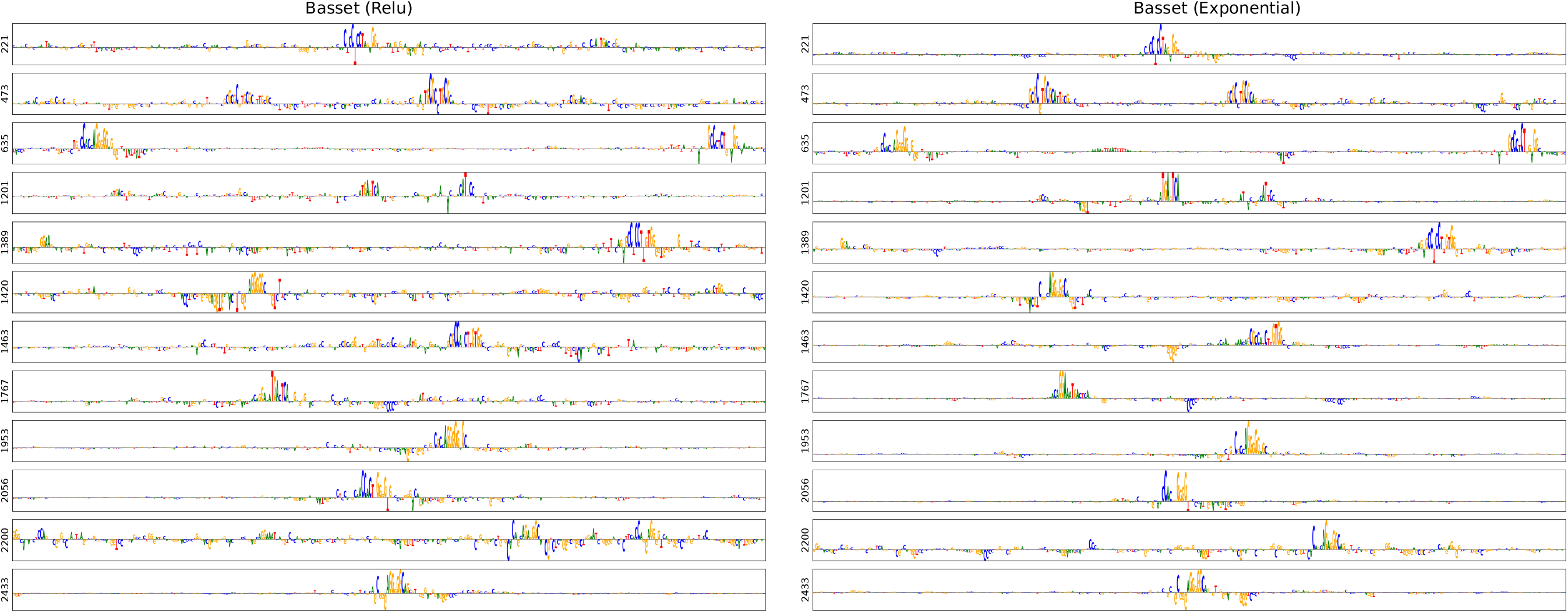
Comparison of attribution scores for sequences about chromatin accessible sites in fibroblast cells. Sequence logo of a saliency map generated for representative test sequences about DNase-seq peaks (Task 4) from a CNN model with relu activations (left) and exponential activations (right). Plots show a 300 nucleotide region centered on each test sequence. The class label in the Basset dataset is 4 (zero-indexed). The *y*-axis labels indicate the sequence index from the test set.

**Supplemental Figure 13.**
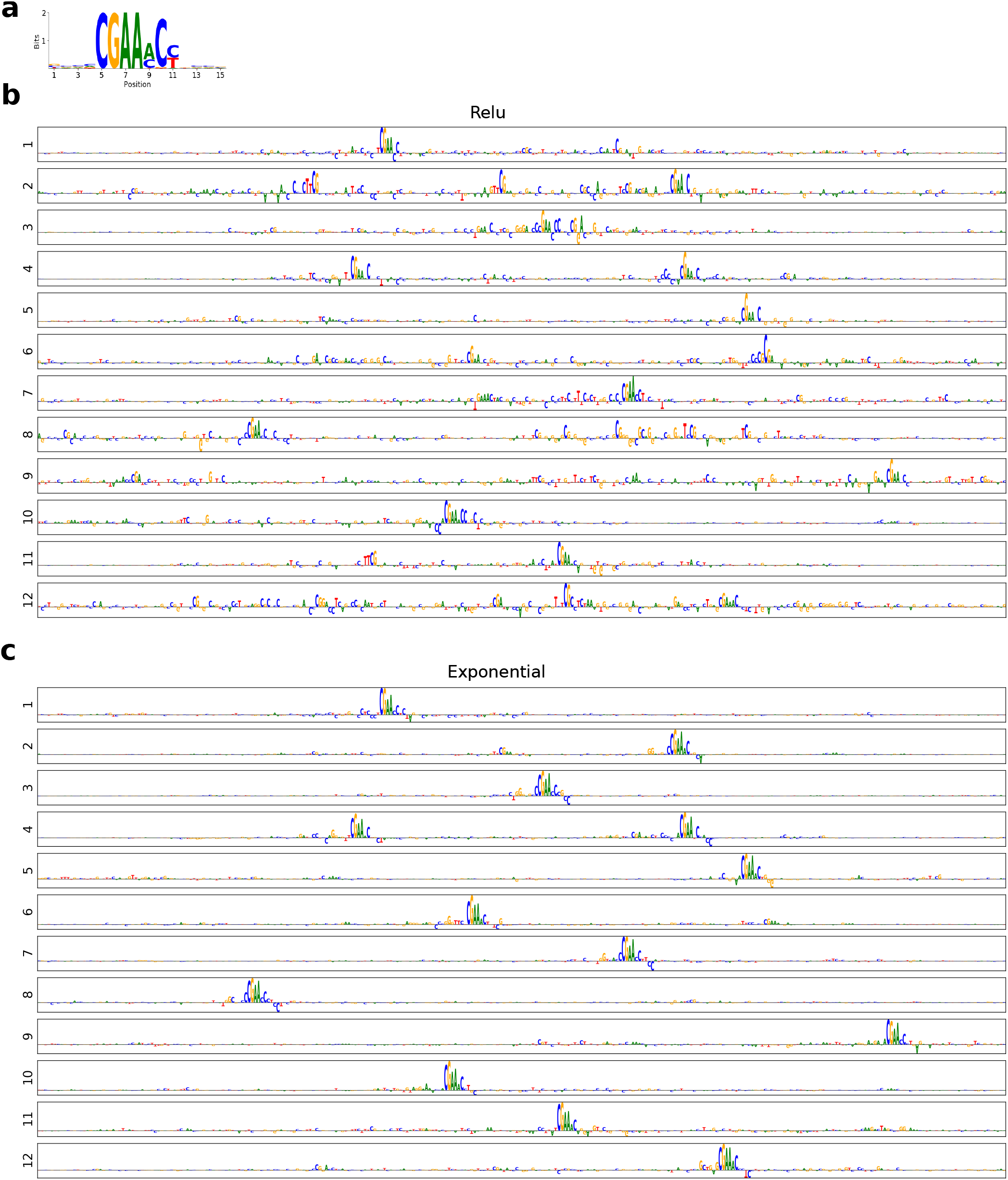
Comparison of attribution scores for sequences about ZBED2 ChIP-seq peaks. (**a**) Sequence logo of the known motif for ZBED2. Sequence logo of a saliency map generated for representative test sequences about ZBED2 ChIP-seq peaks (Task 5) from a CNN model with relu activations (**b**) and exponential activations (**c**).

**Supplemental Figure 14.**
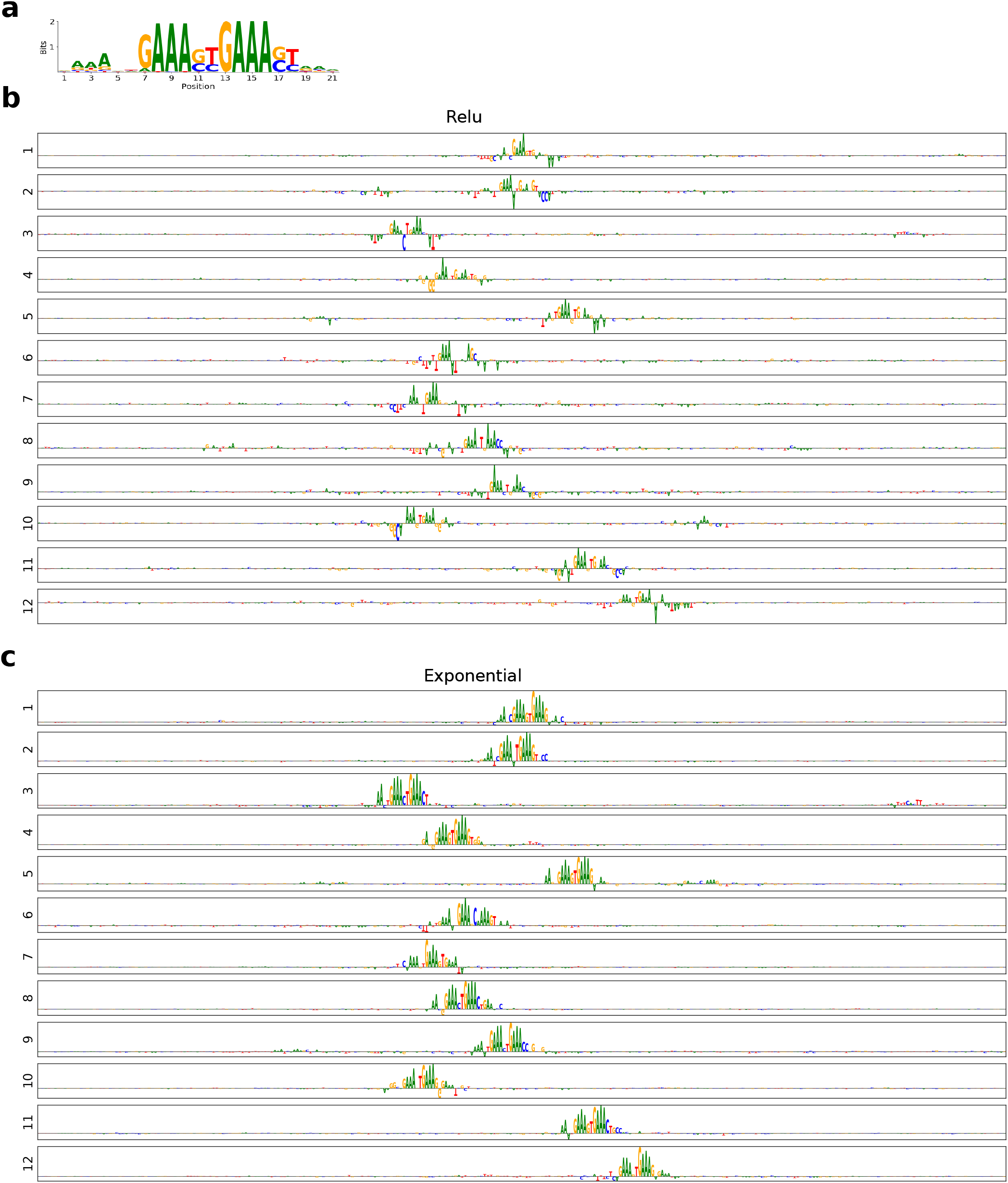
Comparison of attribution scores for sequences about IRF1 ChIP-seq peaks. (**a**) Sequence logo of the known motif for IRF1 from the JASPAR database. Sequence logo of a saliency map generated for representative test sequences about IRF1 ChIP-seq peaks (Task 6) from a CNN model with relu activations (**b**) and exponential activations (**c**).

**Supplemental Figure 15.**
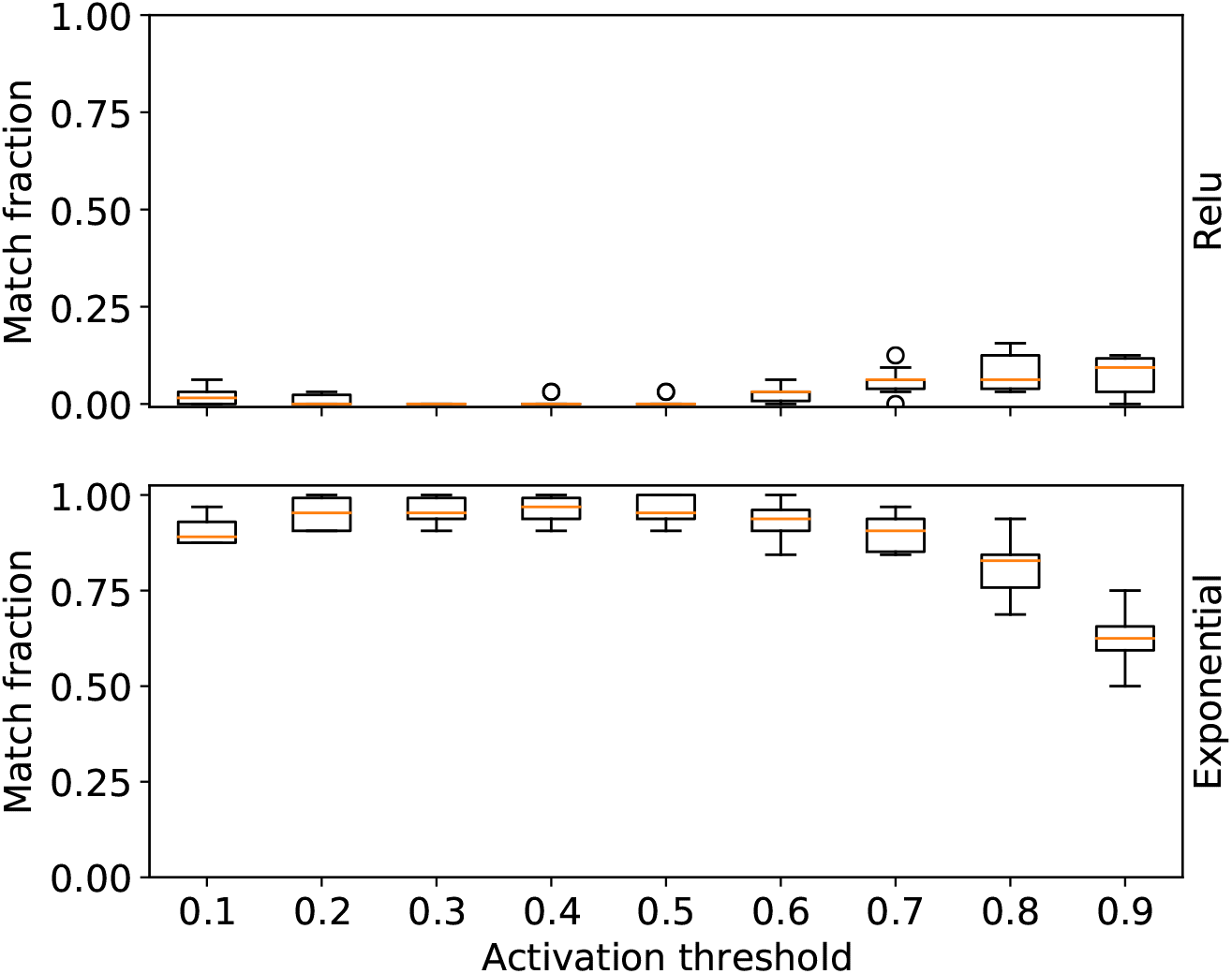
Task 1 filter performance comparison for CNN-deep with different activation thresholds. Box plot of the fraction of filters that match ground truth motifs for CNN-deep with different relu activations (top) and exponential activations (bottom) for different activation thresholds to for the activation-based alignment filter visualization. The default threshold used in the paper is 0.5. Each box plot represents the performance across 10 models trained with different random initializations (box represents first and third quartile and the red line represents the median).

**Supplemental Figure 16.**
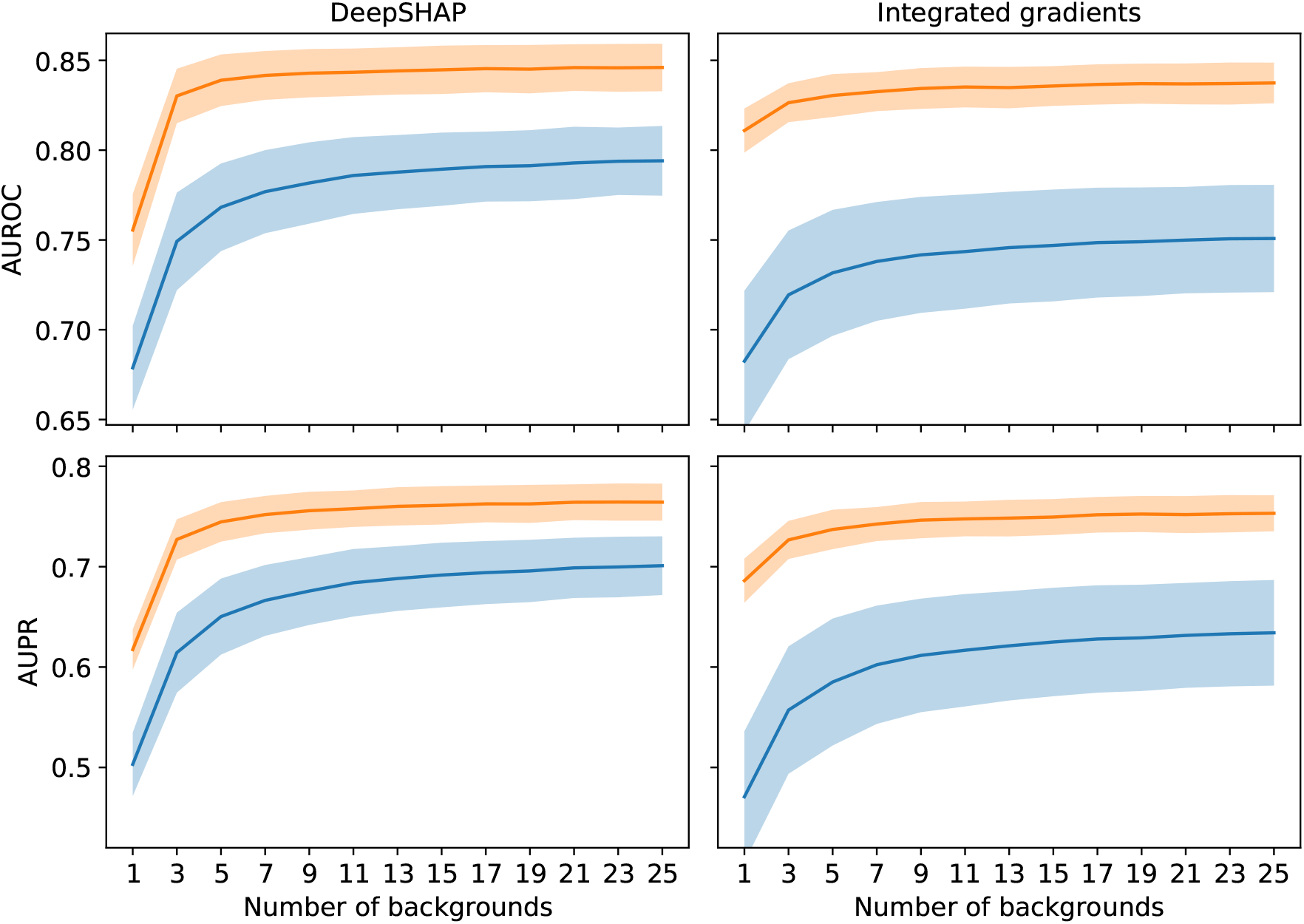
Interpretability performance for attribution methods that rely on reference sequences. Plots of the average interpretability AUROC (top) and AUPR (bottom) for CNN-deep using attribution scores generated on test sequences with DeepSHAP (left) and Integrated gradients (right) for different numbers of randomized reference sequences. The shaded region represents the standard deviation of the mean of the performance across 10 models with different random intializations. The blue and orange lines represent CNN-deep with relu activations and exponential activations, respectively.

